# Mapping Eelgrass Cover and Biomass at Izembek Lagoon, Alaska, Using In-situ Field Data and Sentinel-2 Satellite Imagery

**DOI:** 10.1101/2024.08.07.607047

**Authors:** David C. Douglas, Michael D. Fleming, Vijay P. Patil, David H. Ward

## Abstract

Two eelgrass (*Zostera marina*) maps of Izembek Lagoon, Alaska, were generated by first creating maps of spectrally unique classes from each of two Sentinel-2 satellite images collected on July 1, 2016, and August 14, 2020, then attributing the spectral classes with information about eelgrass conditions based on field data. Maps depicting various eelgrass metrics, such as percent cover and modeled biomass, were generated using summaries of the ground data that spatially intersected each spectral class. Comparisons between the 2016 and 2020 Sentinel-2 maps of eelgrass distributional extent, as well as a 2006 Landsat map, indicated that areas where eelgrass presence may have declined between 2006 and 2020 were most prevalent in the central part Izembek Lagoon, while areas of possible biomass decline were more prevalent in the southern part between 2016 and 2020. Monitoring eelgrass conditions at Izembek Lagoon with satellite imagery and concurrent ground data provides capabilities for making comparisons over time, but the influences of tide levels, growing season phenology, and spatiotemporal co-registration accuracy should be considered when designing and interpreting change detection analyses.

## Background

Izembek Lagoon is a large shallow embayment (∼50 km long x 10 km wide) on the Bering Sea side of the Alaska Peninsula within the Izembek National Wildlife Refuge and Izembek State Game Refuge. Maximum tidal ranges are modest (∼1 m), waters are generally clear, salinity ranges from ∼26 ppt to 32 ppt (McRoy 1966), and the maritime climate is cool with mean monthly air temperatures ranging from -2 to 11 °C annually (Brower et al. 1988).

Storms commonly occur throughout the year and ice cover forms intermittently between December and March (McRoy 1966). The lagoon is dominated by broad tidal flats interspersed with networks of channels stemming from the lagoon’s three entrances at the northern, central, and southern ends. Eelgrass (*Zostera marina*) is the predominant submerged aquatic vegetation. More about the physical and biological environment at Izembek Lagoon can be found in Ward et al. (1997).

Izembek Lagoon supports one of the largest eelgrass beds in the world (McRoy 1966), lending to its designation as a wetland of international importance (Ramsar 2023). Eelgrass provides a key source of nutrients for the lagoon’s food web which includes numerous species of birds, fish, and marine mammals (Ward et al., 1997). Owing to 1) the importance of Izembek Lagoon ecologically, 2) the growing threats to eelgrass along the United States (U.S.) mainland Pacific coast (Sherman and DeBruyckere 2018), and 3) the environmental changes presently occurring in the Bering Sea (Wood et al. 2015, Overland et al. 2024), the U.S. Geological Survey and the U.S. Fish and Wildlife Service have developed a 3-tiered strategy for monitoring eelgrass beds at Izembek that targets different spatial and temporal scales (Neckles et al. 2012, Hogrefe et al. 2014, Ward and Amundson 2019). The broadest-scale monitoring (tier-1) utilizes satellite imagery every ∼5 years to delineate the spatial extent of eelgrass beds throughout the lagoon.

This report describes the most recent implementation of tier-1 eelgrass monitoring at Izembek Lagoon. Two suites of eelgrass maps were created using Sentinel-2 satellite imagery (Phiri et al. 2020) collected in 2016 and 2020. We applied unsupervised (mathematical only) spectral classification techniques to the imagery and then used field data (Ward 2021) to quantitatively attribute the spectral classes with associated eelgrass characteristics (e.g., cover and biomass).

We present the image classification methods first, followed by the methods for characterizing the spectral classes based on field data, and conclude with examples of estimating changes in eelgrass cover and biomass.

Sentinel-2 imagery possess three notable advantages over Landsat: 1) 10-meter resolution as opposed to 30-meter; 2) a revisit frequency of 5 days as opposed to 8 days; and 3) spectral data with higher radiometric resolution (14-bit after scaling) compared to the 8, 12, and 14-bit resolution recorded by Landsat satellites 1-7, 8, and 9, respectively. The 5-day revisit frequency of Sentinel-2 is achieved by two satellites (Sentinel-2A and Sentinel-2B) with 10-day revisit cycles that are maintained in opposing orbits (Li and Roy 2017). Sentinel-2’s polar orbit around the earth is sun synchronous with a late-morning local overpass time. Landsat presently has an 8- day repeat cycle also owing to two operational satellites (Landsat-8 and 9) in opposing 16-day orbits.

To accurately assess eelgrass conditions and changes across years, image acquisition would ideally occur at a low-tide under cloud-free conditions during the peak eelgrass growing season (July and August). Because the probability is low for a satellite overpass to occur under all those conditions at Izembek Lagoon, Sentinel-2’s more frequent revisitation rate compared to Landsat affords a distinct advantage. Further, like Landsat but with up to 9-times the spatial resolution, the Sentinel-2 image swath width (290 km) is sufficient to capture the entirety of Izembek Lagoon in a single pass, so eelgrass mapping is not compromised by having to mosaic imagery from different dates and tide levels. Also, like Landsat (Loveland and Irons 2016), Sentinel-2 collects high-quality radiometric data across a similar range of spectral bands (Drusch et al. 2012).

### Image Acquisition

Sentinel-2 imagery is freely available from the European Space Agency’s (ESA) Copernicus Data Space Ecosystem. The most efficient strategy for searching the Sentinel-2 archive for mapping eelgrass at Izembek Lagoon is to determine which dates between June and September (ideally in July or August when eelgrass abundance peaks) had low-tides (<0.0 MLLW, mean lower low water) during late-morning local time. We used the National Oceanic and Atmospheric Administration (NOAA) Tides and Currents website to acquire tide information for the station at Grant Point, Izembek Lagoon, Alaska (station ID 9463058, https://tidesandcurrents.noaa.gov/noaatidepredictions.html?id=9463058). After we created a list of dates with late-morning low tides, we interrogated a satellite imagery viewing portal like NASA Worldview (https://worldview.earthdata.nasa.gov/) to ascertain the cloud conditions on each late-morning low-tide date. We found most dates to be cloudy, but a few low-tide dates with complete or partial visibility of Izembek Lagoon were advanced to the next step of checking the Sentinel-2 archive to determine if: 1) a Sentinel-2 image was collected on the respective date, and if so 2) does the image quality appear acceptable to meet the eelgrass mapping objectives.

For mapping eelgrass, we specifically downloaded the Sentinel-2 Level-1C product which provided top-of-atmosphere (TOA) reflectances in cartographic geometry. The Sentinel-2 imagery is distributed as UTM/WGS84-projected tiles of approximately 110 x 110 km dimension on a 100 km grid, so there is ample overlap with adjacent tiles if necessary. Sentinel-2 products are comprised of 13 bandwidths of spectral wavelengths: four bands at 10-meter resolution, six bands at 20 m and three bands at 60 m (Figure 1).

**Figure 1.**
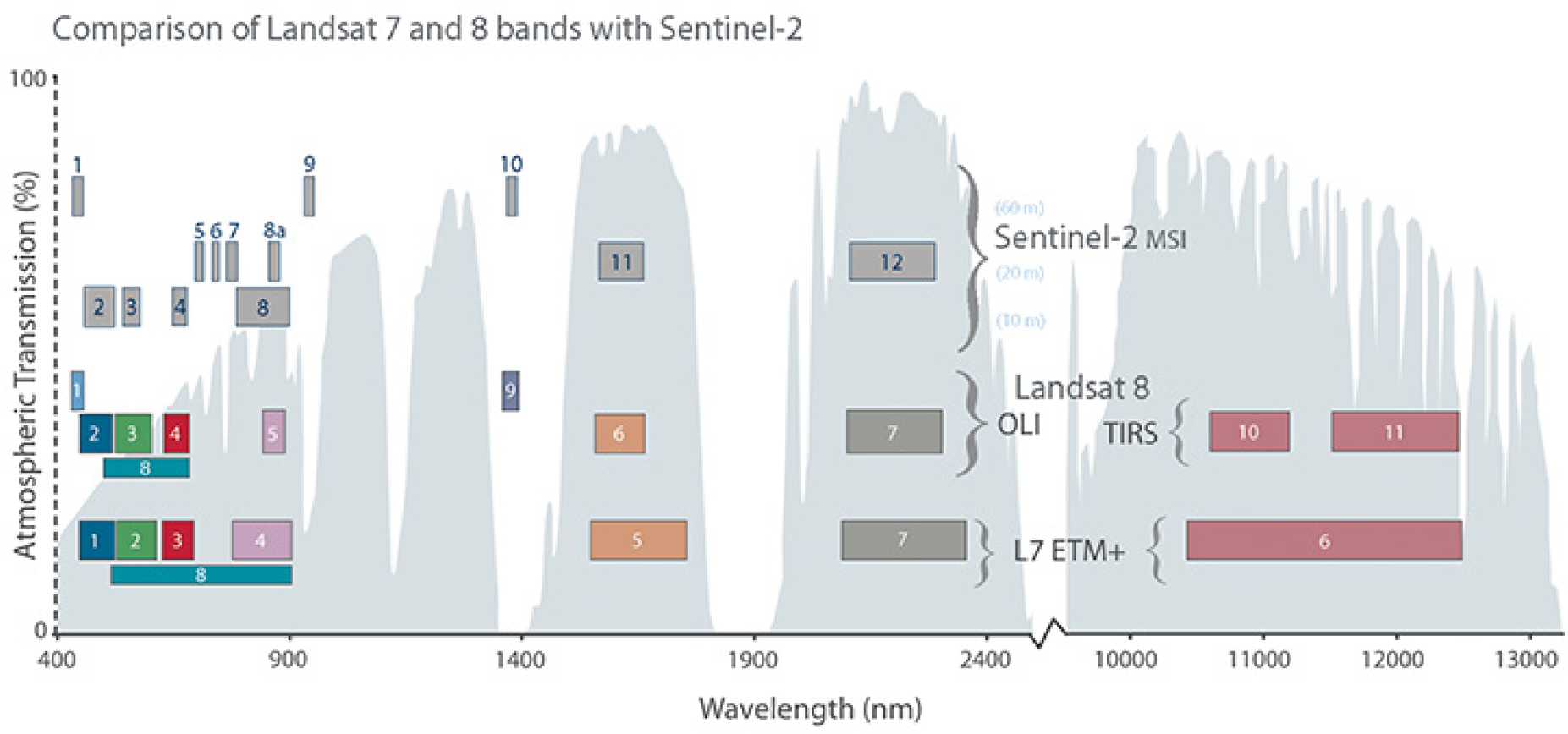
Comparison of Landsat 7 and Landsat 8 spectral bands with Sentinel-2 bands. https://landsat.gsfc.nasa.gov/wp-content/uploads/2015/06/Landsat.v.Sentinel-2.png (public domain).

Our searches of the Sentinel-2 archive from 2008 to 2023 during the eelgrass growing season found two dates when clear-sky, low-tide, and image-acquisition all coincided, thus providing data suitable for spectral classification and eelgrass mapping throughout Izembek Lagoon (Table 1).

**Table 1.** All Sentinel-2 archived imagery suitable for mapping eelgrass at Izembek Lagoon, Alaska, during a 16-year period (2008–2023). The Level-1C archival image granule (tile) names are documented below the acquisition dates. Tide levels (feet) relative to mean lower low water (MLLW) are shown at the time of image collection. The years of ground data (Ward 2021) that contributed to map development and attribution are also shown.

| Image acquisition date (UTC) | Satellite | Tide (ft, MLLW) | Ground data years |
| --- | --- | --- | --- |
| 2016-07-01 22:05:32<br>(L1C_T03UXB_A005360_20160701T220836) | Sentinel-2A | -0.7 | 2015, 2016 |
| 2020-08-14 22:30:31<br>(L1C_T03UXB_A017973_20200814T220533) | Sentinel-2B | -0.1 | 2018, 2019, 2022 |

### Image Pre-Processing

We downloaded the Sentinel Level-1C image tiles containing Izembek Lagoon for each of the two dates shown in Table 1. The Level-1C product was delivered as terrain corrected top-of-atmosphere reflectances (the ratio between the incoming light and the light reflected after travelling through the atmosphere) in a georeferenced format (UTM Zone 3N), but otherwise the Level-1C had the fewest alterations applied to the raw satellite image swath data compared to other Sentinel-2 image data formats. Each single tile from the 2 dates was cropped to the same 5000-line by 6000-sample image subset that fully encompassed Izembek Lagoon with 10-meter pixel resolution and an upper left origin-pixel center-coordinate of ULX=6100005 and ULY=6155995. We resampled the 20-meter bands to10-meter resolution using bilinear interpolation and stacked all bands into a 10-band composite image (Table 2) for subsequent spectral analysis.

**Table 2.** Structure and contents of a Sentinel-2 10-band composite image used for spectral analysis. Resolutions and center wavelengths of each band are shown. Band wavelengths are generalized into 6 categories: B=blue, G=green, R=red, NIR=near infrared, VNIR=very near infrared, and SWIR=shortwave infrared. Spectral bandwidths are illustrated in Figure 1.

| Composite band # | 1 | 2 | 3 | 4 | 5 | 6 | 7 | 8 | 9 | 10 |
| --- | --- | --- | --- | --- | --- | --- | --- | --- | --- | --- |
| Sentinel-2: |  |  |  |  |  |  |  |  |  |  |
| band # | 2 | 3 | 4 | 8 | 5 | 6 | 7 | 8A | 11 | 12 |
| resolution (m) | 10 | 10 | 10 | 10 | 20 | 20 | 20 | 20 | 20 | 20 |
| wavelength (nm) | 490 | 560 | 665 | 842 | 705 | 740 | 783 | 865 | 1610 | 2190 |
| wavelength category | B | G | R | NIR | VNIR | VNIR | VNIR | NIR | SWIR | SWIR |

We originally attempted to identify areas of change by clustering a combined 2-date, 20-band image into 80 classes using the unsupervised algorithm ISODATA in the open-source software package MultiSpec© (version 9.2011, https://engineering.purdue.edu/~biehl/MultiSpec/). The class statistics were used with a Gaussian maximum likelihood classifier (MLC) to assign each image pixel membership into one of the cluster classes. Evaluation of that classification, however, showed that the classes most often associated with change were dominated by changes in the physical environment such as alterations in mudflats, high tide areas, and deep-water channels as opposed to changes in the biological environment. Consequently, we only used the combined 2-date classification to isolate the lagoon area by grouping all classes within the lagoon (except for obvious deep-water classes) and masking all remaining classes (i.e., upland vegetation and deep water). An elevation mask (elevation >0 m) was additionally applied to exclude onshore water. The resulting mask included the intertidal areas of Izembek Lagoon (not areas of deep water), and some neighboring embayments (i.e., Cold Bay and Kinzarof Lagoon) that were within the footprint of the Sentinel-2 image subsection. This mask was applied to both 10-band sub-sectioned images (i.e., 2016 and 2020), and thereafter the two masked images were analyzed independently.

### Image Spectral Analysis

Image analysis involved two parts. First, splitting the image pixels into cluster classes, then second, grouping the cluster classes into spectral classes.

The splitting phase used ISODATA and MLC to derive and map 40 cluster classes from each 10- band masked image, hence the cluster classes were specific to Izembek Lagoon proper. Forty clusters were prescribed — roughly twice the desired number of final classes.

The second phase grouped the cluster classes into a set of classes that were homogenous both spectrally and spatially. A series of iterations were performed to group the cluster classes into spectral classes. The overarching strategy was to maximize the number of spectral classes while reducing duplication of very similar classes.

We applied five strategies to aid in the grouping process:

1. Calculation of a saturating transformed divergence metric, a measure of spectral separability between classes (Swain and King 1973).
2. Calculation of an occurrence matrix that indicates which cluster classes are next to other classes spatially.
3. A chaining algorithm that produces hierarchical cluster relationships run on their separability (#1 above), their occurrence (#2), and on both combined (Aucoin and Stewart 1978).
4. Assessment of spectral biplots that graphed cluster means for 2 bands (i.e., SWIR x NIR, Red vs NIR, etc.; for example, Figure 2).
5. Calculation of the number of ground data points in each cluster class to ensure the final spectral classes had a reasonable sample of field data for characterizing the associated eelgrass conditions.

**Figure 2.**
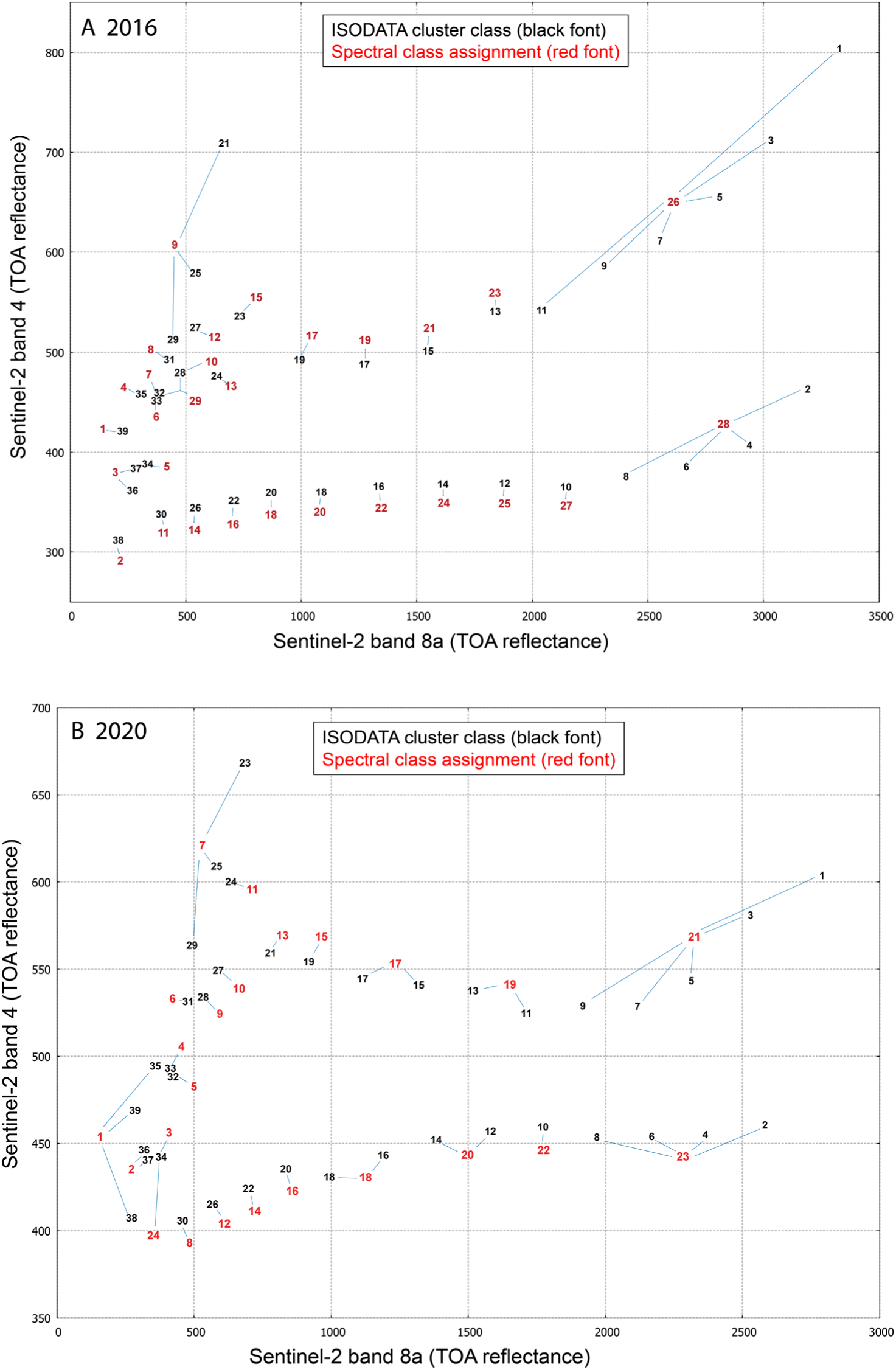
Spectral biplots of cluster class means (denoted with black font) plotted on top of atmosphere (TOA) reflectances (unitless) of two spectral bands. Cluster class membership in grouped (or split) spectral classes are denoted with red font (and positioned for legibility) for the (A) 2016 and (B) 2020 Sentinel-2 classifications of Izembek Lagoon, Alaska. This figure illustrates one of several strategies that guided the manual assignment of cluster classes into spectral classes, as summarized in Table 3.

**Table 3.**
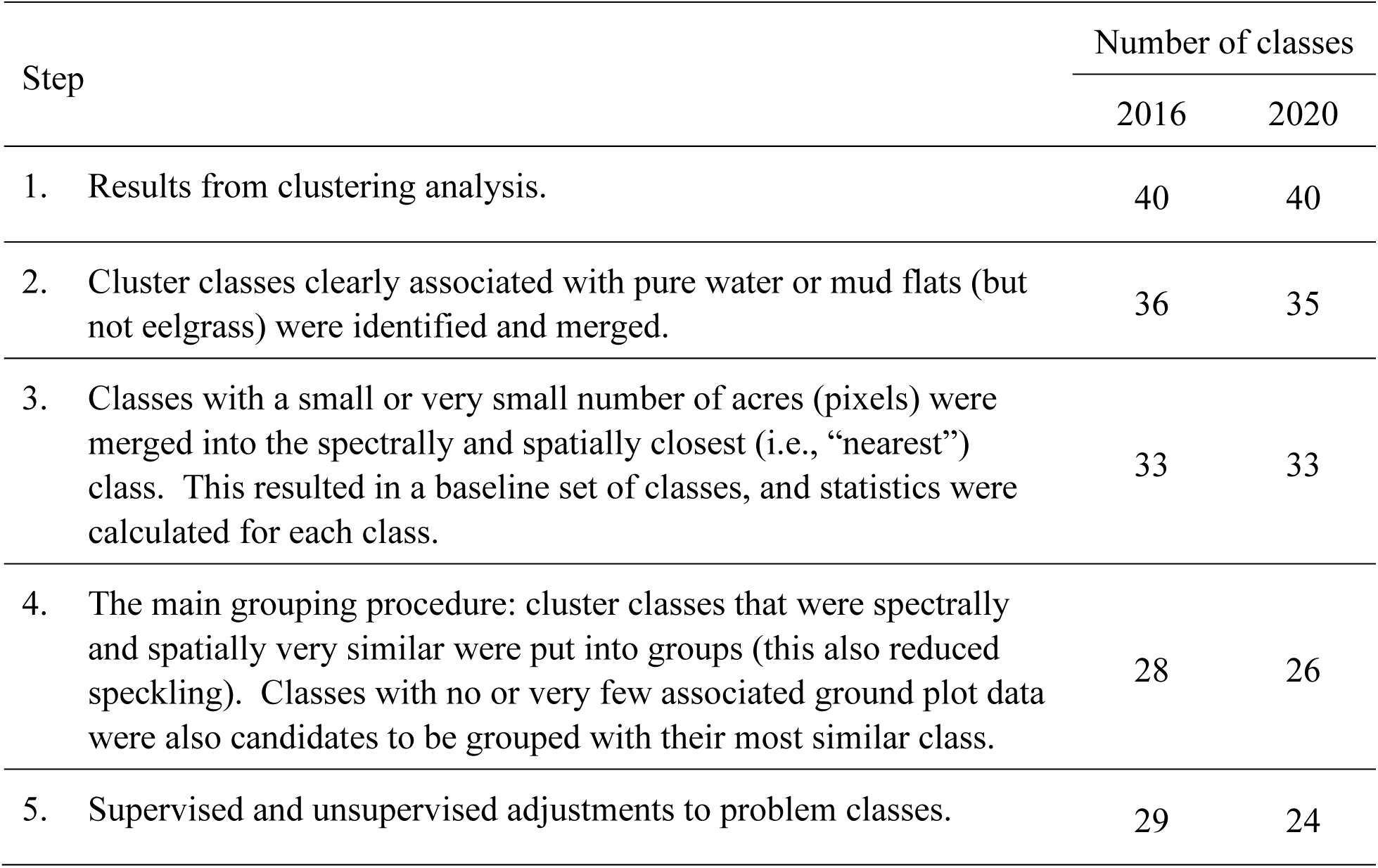
Sequence of steps applied to assign 40 cluster classes to spectrally and spatially similar spectral classes for each of the two Sentinel-2 image analyses (2016 and 2020) for Izembek Lagoon, Alaska.

These strategies were collectively used to guide a visual and numerical interpretation of the cluster classes with the primary goal of grouping similar classes. First, cluster classes clearly associated with pure water or mud flats (but not eelgrass classes) were identified and merged. This merging resulted in 36 and 35 cluster classes for the 2016 and 2020 images, respectively. Next, classes with a small or very small number of pixels were grouped into the spectrally and spatially closest (i.e., “nearest”) class. Then, cluster classes that were spectrally and spatially very similar were grouped. Last, classes with no or very few ground data were grouped with their most similar class. These steps served to create spectrally and spatially homogeneous spectral classes by aggregating similar entities from the clustering step and ensuring each class had field data. Figure 2 illustrates the assignment of cluster classes to spectral classes, mostly by grouping but with a few instances of splitting and masking. The biplots in Figure 2 show an example of spectral relationships between just two of Sentinel’s 10 spectral bands, meaning Figure 2 shows an incomplete representation of the many spectral relationships among clusters. Figure 2 is presented here simply to illustrate how biplots can be a useful tool for guiding cluster groupings.

Following critical inspection of the spectral classes, two types of targeted adjustments to “problem classes” were applied for purposes of attaining better spectral continuity. One type involved splitting a spectral class and the other involved masking pixels for supervised class reassignment. Specifically, in the 2020 image, one cluster appeared to contain at least two different classes based on its spectral and spatial characteristics. That cluster was split into 10 classes using the 10-band spectral data and the ISODATA clustering algorithm. The 10 classes were assessed spectrally and spatially and grouped into 2 classes resulting in one class occurring primarily in the southernmost part of the lagoon and appearing to be associated with deep water, and the other class more in the northern lagoon associated with shallower water. The 2020 clustering also required a supervised reassignment to the classification. A few isolated cloud shadows were present on the mud flats in the northeastern lagoon, so a mask boundary was digitized by hand to encompass each area of cloud and cloud shadow, then the cluster classes within the cloud mask (in this case two classes) were changed to match the class of the surrounding mud flats. A similar supervised fix was applied to the 2016 map that involved remapping affected classes to a surrounding mudflat class. The last adjustment to both maps targeted areas on mud flats where large drifting accumulations of dislodged eelgrass and green seaweed often get deposited by tidal actions, and not surprisingly get included in spectral classes associated with eelgrass. Using expert knowledge (D. Ward), we hand digitized a custom mask of areas where large entanglements of drifting eelgrass and green seaweed often wash up onto mud flats, then recoded eelgrass-associated spectral classes within that mask to a mud flat class.

After all the groupings and custom adjustments described above were applied, there were 29 spectral classes for the 2016 Sentinel-2 image and 24 classes for the 2020 image. The sequence of steps and outcomes described above are summarized in Table 3.

### Field Data Preparation

Field data (Ward 2021) that were collected on-site at spatially coincident sample plots were used to attribute each respective spectral class with information about the ground cover characteristics (i.e., surface types like eelgrass, mud, water, etc.). Many years of field data have been collected at Izembek Lagoon (Ward 2021) to monitor changes in eelgrass abundance and distribution (2007–2011, 2015–2019, 2022–2023). Recognizing that changes are intrinsic to eelgrass communities (Ward et al. 2003, Bartenfelder et. al. 2022, Munsch et al. 2023), using ground data collected within a year or two of a satellite image collection date reduces the chances of the field data misrepresenting the actual ground conditions as recorded by the satellite image. For the 2016 Sentinel-2 image, we used field data collected in 2015 and 2016, and for the 2020 Sentinel-2 image, we used field data collected in 2018, 2019 and 2022 (Table 1). We also added ground data from plots sampled in 2007 and 2008 where no eelgrass was observed, but only if those plots were not resampled in subsequent years because they had since remained in high intertidal areas beyond the growth range of eelgrass in the lagoon (Ward et al. 1997). Including the 2007 and 2008 ground data collected at persistent mudflats bolstered sample sizes for characterizing their associated spectral classes.

Data at each field sampling plot were recorded at four 0.25 m^2^ quadrats, arbitrarily positioned diagonally 5-10 meters from the plot location in each of four compass-oriented quadrants (NW, SW, NE, SE; Ward 2021). We diagonally positioned the UTM location of each sampled quadrat by adding and/or subtracting 10 m to the plot’s UTM easting and northing, respective to the quadrant’s diagonal (NW, SW, NE, SE) compass direction. Including each of the four sampled quadrants at each plot, as well as the 2007–2008 unvegetated plots, bolstered sample sizes for characterizing the spectral classes with information about the associated ground conditions.

Four variables recorded at each plot quadrat were specifically relevant for quantifying eelgrass conditions associated with each spectral class (Table 4, percent cover, Braun-Blanquet cover category (Braun-Blanquet 1972), presence, and abundance index). If a plot was sampled in more than one year, averages were calculated for each quadrant subplot.

**Table 4.** Variables from the field data (Ward 2021) that were used to quantify characteristics of each spectral class in terms of eelgrass fractional cover, presence, abundance, and biomass at Izembek Lagoon, Alaska.

| Variable name | Variable description |
| --- | --- |
| Zm_percent | Percent cover of eelgrass in the 0.25 m<sup>2</sup> quadrat; percentages recorded as <1.0 denote: 0= no shoots; 0.1= 1 shoot present; 0.5= 2-5 shoots present. |
| Zm_BB | Braun-Blanquet (BB) visual estimation category of eelgrass cover: 0=0% cover; 0.1=1 shoot; 0.5=2 shoots to 4% cover; 1.0-1.8 = 5-20% cover; 2.0-2.8=25-45% cover; 3.0-3.8=50-70% cover; 4.0-4.8=75-95% cover; 5.0=100% cover. |
| Zm_pres | Eelgrass was present (1) or was not present (0) in the sample quadrat. |
| Zm_abunIndex | Abundance index of eelgrass. Calculated as the mean shoot length (mm) multiplied by Zm_BB |
| Zm_biomass | Model-estimate of eelgrass biomass based in part on data from destructive collection of plot samples and measured as dry weights after desiccation. Biomass was estimated at each plot using a Bayesian gamma hurdle model which models the probability of occurrence and eelgrass abundance (biomass) where present as separate processes (Patil 2024). More information about the model can be found in Ward and Amundson (2019). |

### Spectral Class Attribution

The ground-plot quadrat data were spatially intersected with the spectral classes to calculate class-specific eelgrass cover statistics (mean, standard deviation, median, and interquartile range). It was not uncommon for some spectral classes to be associated with notably different characterizations of ground cover. There are several explanations why a spectral class could be characterized by a breadth of ground data conditions, including: 1) real changes in ground conditions that occurred between the dates of field sampling and satellite image collection; 2) accuracy of the field plot locations; 3) co-registration accuracies between the field plot locations and the georeferenced satellite image; 4) representativeness of the 0.25 m^2^ quadrat relative to the 10 m x 10 m Sentinel-2 grid cell; and 5) the choice of which Sentinel-2 grid cells to assign to each of the four sampled quadrants at each plot. For the latter, we assigned the sampled quadrants to the four grid cells on the diagonals of the grid cell that intersected the plot’s center location as opposed to the four cells in the cardinal directions.

Recognizing the high likelihood that some misrepresentative ground data gets included in calculations of class-specific eelgrass cover statistics, before those statistics were finalized, we first excluded the more egregious outliers for each variable (Table 4) within each spectral class. Outliers were defined to be any data records with values outside (above or below) 1.5*IQR (IQR=interquartile range). After the outliers were excluded, we recalculated and graphed the class-specific statistics for each eelgrass variable as violin plots (Figures A1-A4) and tabled the corresponding means, medians, and variances (Tables A1-A8). The modeled eelgrass biomass estimates, and 95% credible intervals (Patil 2024) were calculated for each field plot location (not quad) so sample sizes intersecting each spectral classes were smaller (Figure A5).

Consequently, we chose to derive biomass maps based on median values only (Tables A9, A10) because the mean values were sometimes unrealistically skewed.

### Eelgrass Mapping

Our eelgrass mapping strategy at Izembek Lagoon began with unsupervised derivation of statistical cluster classes which we subsequently grouped into spectral classes, and then attributed with environmental data from field studies. In other words, the spectral classes established a gradient (or scale) of environmental conditions and the ground data served to calibrate that scale. The underlying assumption of this strategy is that surface areas with similar cover characteristics will possess similar spectral characteristics, which in turn will be grouped (clustered) into like spectral classes, and thus provide useful information about the spatial distributions of surface cover types throughout the study area.

Eelgrass maps were produced by attributing each spectral classes with an eelgrass statistic of choice derived from field observations (after outliers had been excluded). Maps were generated for each of the five eelgrass variables in Table 4. For example, a map of median eelgrass percent cover was produced by deriving that statistic for each spectral class from the ground plot data that spatially intersected the respective classes (Figure 3 A,B), and similarly, a map of median eelgrass biomass was produced (Figure 3 C,D).

**Figure 3.**
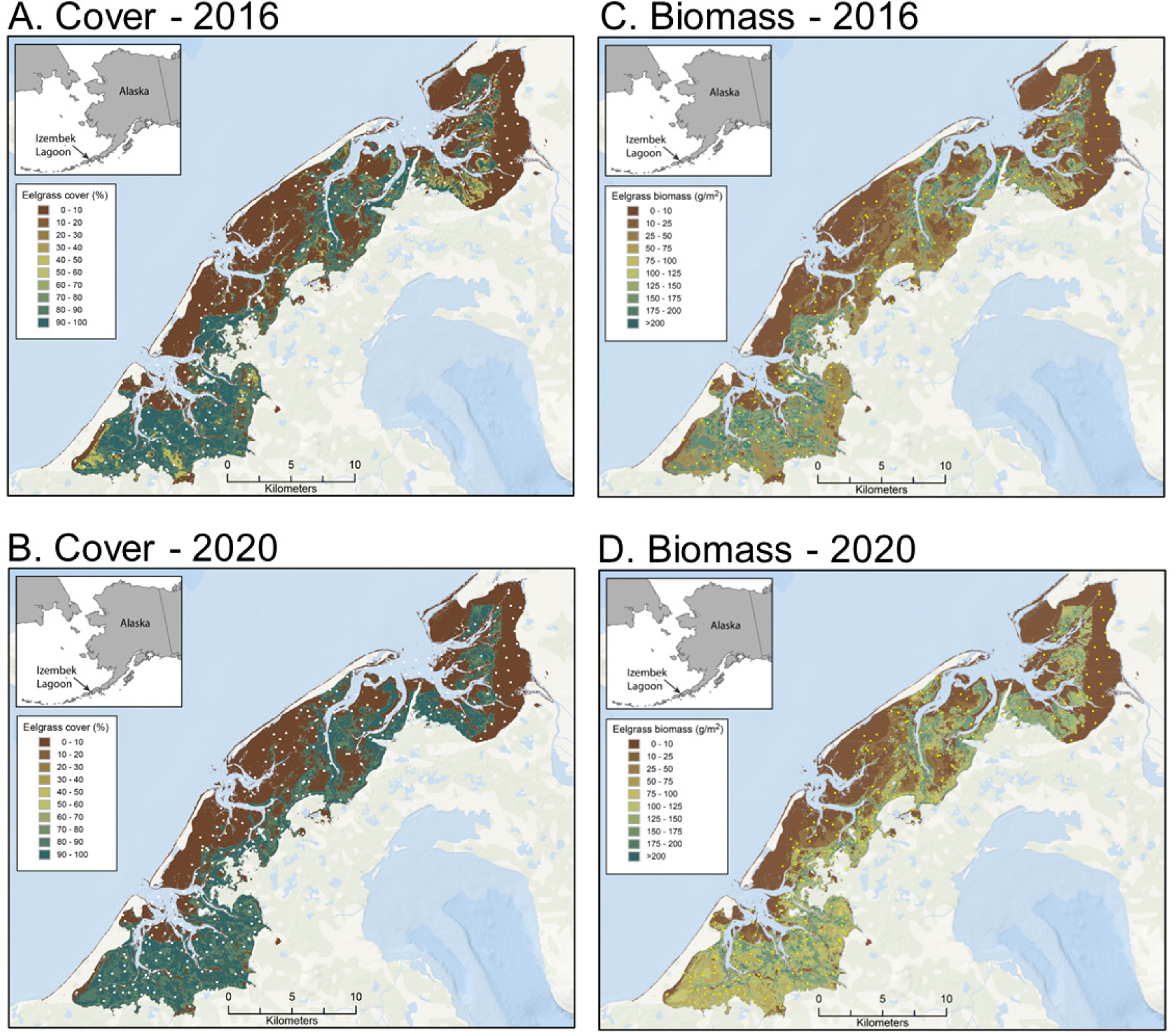
Maps of median eelgrass cover (A,B) and eelgrass biomass (C,D) at Izembek Lagoon, Alaska, derived from Sentinel-2 satellite images collected during low tide on July 1, 2016 (A,C) and August 14, 2020 (B,D). Twenty-nine [24] spectral classes were generated from the 2016 [2020] Sentinel-2 multispectral images within the lagoon, then field data (Ward 2021, Patil 2024) collected throughout the lagoon were used to characterize each spectral class with respect to the field data collected there. In these maps, each spectral class has been color-shaded based on the median eelgrass as directly observed (cover) or model-estimated (biomass) at spatially coincident field plots (dots).

### Earlier Landsat-derived Eelgrass Maps

Two prior eelgrass maps of Izembek Lagoon were produced from Landsat satellite imagery (Ward et al. 1997; Hogrefe et al. 2014). Both Landsat maps employed ISODATA methods initially, like we did for the Sentinel imagery described above, but thereafter the methods of eelgrass mapping differed and are described below for comparison.

The first eelgrass map of Izembek Lagoon (Ward et al. 1997) was produced from a July 28, 1978, Landsat-3 multispectral scanner satellite image comprised of 4 spectral bands with 50 m pixel resolution collected during a late-morning (11:14 local) tide at -0.3 ft. ISODATA was applied to partition the image data encompassing the lagoon into 33 spectral clusters, then each pixel in the image was assigned to a spectral class using an MLE classifier. Each spectral class was subsequently labelled as water, eelgrass, unvegetated, or upland based on personal knowledge of the area, a review of black and white aerial photographs taken in June 1976, and ground data collected in 1986. Two final steps included changing pixels classified as eelgrass in upland areas to the upland class, and similarly, small areas of eelgrass (≤2 pixels) surrounded by unvegetated areas were changed to unvegetated.

The second eelgrass map of Izembek Lagoon (Hogrefe et al. 2014) was produced from a combination of two Landsat images: one on August 2, 2002, by Landsat-7-ETM+ at a tide level of +0.03 feet and the other on July 20, 2006, by Landsat-5-TM at a -0.27 tide. Two images were used to overcome some isolated cloud contaminations. Each image was classified independently by first masking dry land and clouds and then using ISODATA to derive 35 spectral clusters.

Clusters were assigned to one of eight substrate classes to create eight training regions: four for eelgrass and two each for bare ground and water (or left unassigned if uncertain). Clusters spatially coinciding with areas of presumed eelgrass based on false-color enhancements of Landsat bands 4, 5, and 1 were manually assigned to one of four eelgrass classes based on the relative strengths of band-4 (near infrared) and band-3 (red) radiances: 1) high NIR, 2) moderate NIR, 3) low NIR, and 4) minimal NIR. Similarly, spectral clusters representing bare ground or water were determined by visual interpretation and partitioned into either of two respective subclasses, also based on spectral radiance profiles: dry or wet bare ground, and shallow/turbid or deep/clear water. Then, all assigned clusters were merged to create discrete training clusters for a MLE classification that assigned every pixel to one of the eight substrate classes. To overcome small amounts of cloud contamination, the two maps were mosaiced with the 2002 map covering the southern third of the Lagoon and the 2006 map covering the northern two-thirds. A final manual step changed small, isolated patches of eelgrass on bare ground to a ninth map class, seaweed. We refer to this map henceforth as the 2006 Landsat map.

The initial unsupervised derivation of roughly 35-40 spectral clusters using ISODATA was common to all satellite mapping approaches, Landsat and Sentinel. Thereafter, however, both Landsat approaches employed a greater degree of subjective supervision than did the Sentinel approach. For the 1978 Landsat map, the initial suite of 33 spectral clusters were each manually assigned to one of four map classes based on expert interpretations of aerial photographs, spatial context, and field data. For the 2006 Landsat map, the initial suite of 35 spectral clusters were manually assigned to one of eight training classes based on spatial context as well as spectral relationships among the red and infrared imagery bands. Clusters assigned to each of the eight training classes were then merged, and those eight merged clusters were used with MLE to produce a map specifically of those eight defined substrate classes.

The Sentinel method began by developing a set of spectral clusters with ISODATA, then some were split or merged based on spectral relationships, spatial juxtapositions, and sample sizes. Few presumptions or supervised attributions about surface cover were imposed. Rather, the set of clusters was used to produce a map of spectral classes, then each spectral class was attributed with surface cover information by quantifying ground data collected within each class’s spatial footprint. In other words, our Sentinel mapping method transformed a static map of derived spectral classes into a map of surface cover by attributing each spectral class with a summary of ground data about a metric of interest (for example, Figure 3). With an adequate representation of ground conditions to robustly characterize the spectral classes, this strategy provides objectivity and utility owing to the breadth of map themes that can be derived for a variety of mapping goals.

### Eelgrass Change Detection

Eelgrass changes at Izembek Lagoon can be estimated and quantified by differencing (subtracting) two eelgrass maps depicting conditions in different years. The two maps must represent the same eelgrass metric, so the areas where the two maps disagree should presumably show areas where eelgrass had increased or decreased. However, map-detected changes are susceptible to false positives caused by factors other than real on-the-ground changes. Two influential factors stem from differences in: 1) the precise tide level at the time of satellite image acquisition; and 2) the seasonal phenology of eelgrass growth on the date of image acquisition. Areas of eelgrass that are partially submerged at low tide and/or possess only sparse eelgrass cover will be more challenging to robustly detect in imagery that is collected at a time when the tide is only slightly negative. Similarly, analyzing imagery that is many weeks from the timing of peak eelgrass growth (early August) will tend to diminish eelgrass detectability. These factors (tide levels and phenology) should be considered when interpreting map-derived changes, especially changes involving areas associated with marginal (sparse) eelgrass cover.

The 2006 Landsat map of eelgrass at Izembek Lagoon delineated four spectral classes associated with areas of eelgrass with varying levels of near infrared reflectance (Hogrefe et al. 2014). For purposes of detecting eelgrass changes between the 2006 Landsat map of eelgrass and either the 2016 or 2020 Sentinel-2 eelgrass maps, we considered all four Landsat eelgrass classes to be representative of eelgrass presence. For the two Sentinel-2 eelgrass maps, we considered all spectral classes with a median Braun-Blanquet cover value >1.0 (Figure A6) to be comparable to the four combined Landsat classes of eelgrass. With those definitions, eelgrass presence/absence maps were generated for each of the three mapping years (2006, 2016, and 2020). Each pair of these derivative 2-class (present/absent) eelgrass maps was then differenced to spatially depict where eelgrass was gained, lost, or unchanged from the earlier map year to the later map year.

For illustration, we show results for the three difference maps (Figure 4), as well as the net areal extent (km^2^) of those mapped changes in eelgrass presence (Table 5). Roughly 40 km^2^ of area that supported eelgrass in 2006 was lost by 2020, while about 15 km^2^ of eelgrass was gained, resulting in a net loss of about 25 km^2^ (Table 5). Most areas where the presence of eelgrass was lost between 2006 and 2020 were in the central part of Izembek Lagoon (Figure 4).

**Figure 4.**
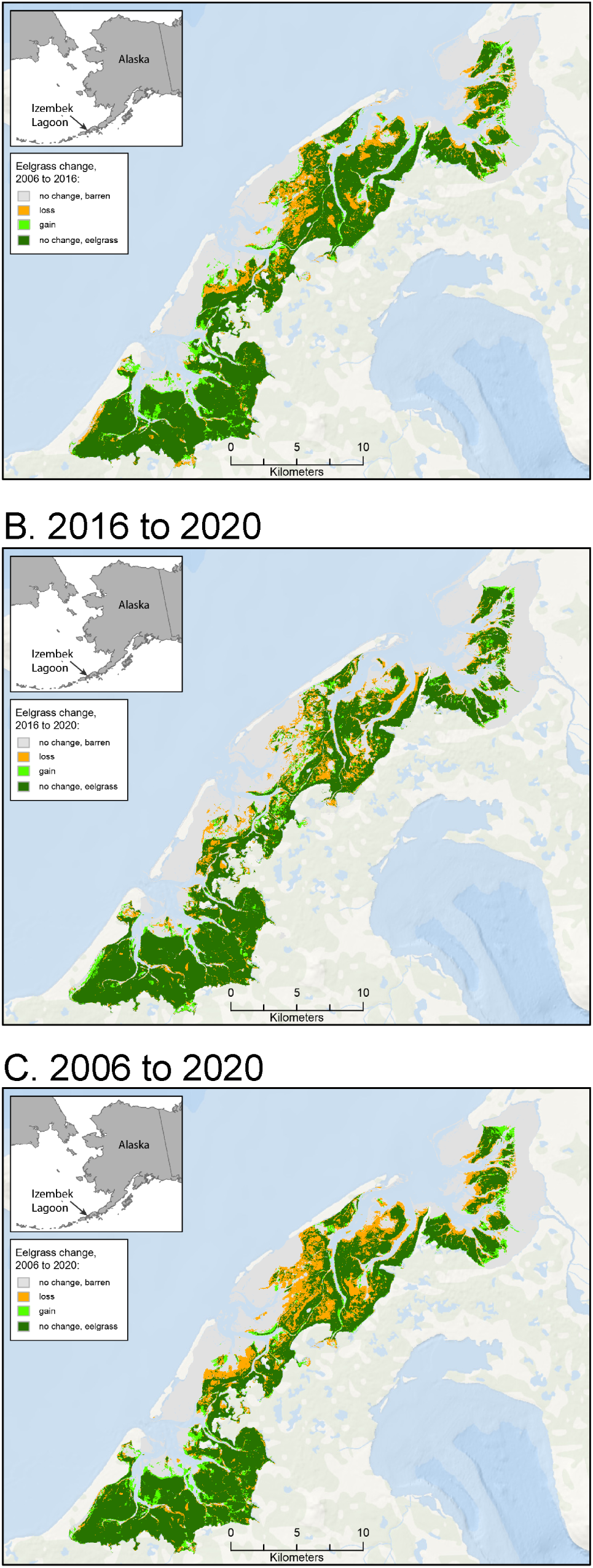
Areas where the presence of eelgrass was gained, lost, or unchanged in Izembek Lagoon, Alaska, from 2006 to 2016 (A), 2016 to 2020 (B), and 2006 to 2020 (C).

**Table 5:**
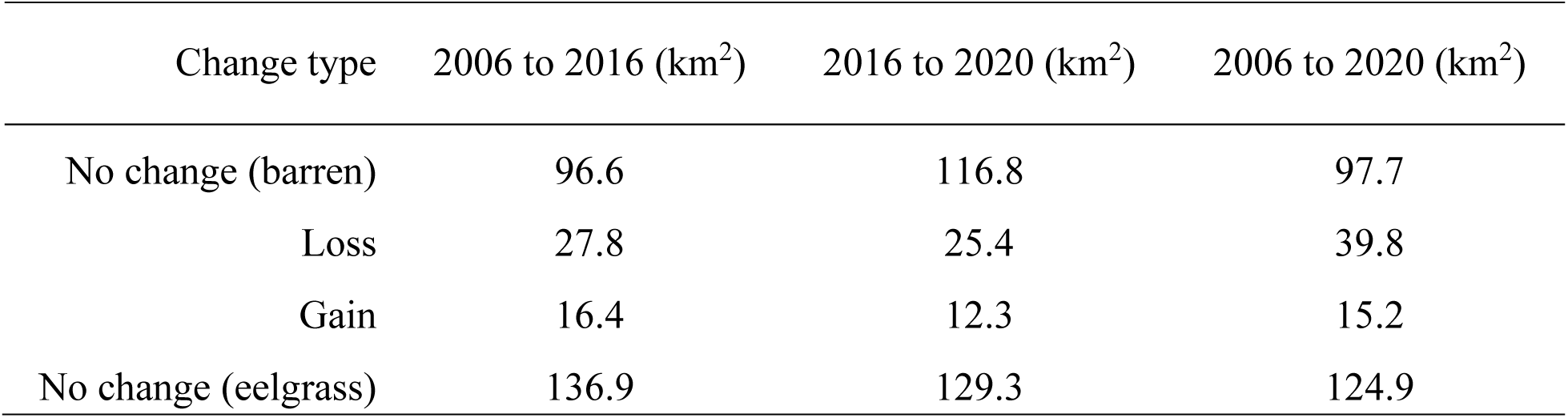
Net spatial extent (km^2^) of eelgrass presence gained, lost, or unchanged in Izembek Lagoon, Alaska from 2006 to 2016, 2016 to 2020, and 2006 to 2020. Area calculations do not include the Lagoon’s deep-water channels.

| Change type | 2006 to 2016 (km<sup>2</sup>) | 2016 to 2020 (km<sup>2</sup>) | 2006 to 2020 (km<sup>2</sup>) |
| --- | --- | --- | --- |
| No change (barren) | 96.6 | 116.8 | 97.7 |
| Loss | 27.8 | 25.4 | 39.8 |
| Gain | 16.4 | 12.3 | 15.2 |
| No change (eelgrass) | 136.9 | 129.3 | 124.9 |

Spatial changes in estimated modeled biomass from the first Sentinel-2 imagery date (July 1, 2016) to second imagery date (August 14, 2020) are shown in Figure 5 and were calculated by subtracting the two biomass maps presented in Figure 3. While eelgrass biomass changes significantly over the course of a growing season (Ward et al. 2022), the biomass model adjusted for day of year and tide level (Ward and Amundson 2019), so the two maps were better standardized with respect to phenology and thus better suited for revealing between-year changes in community density or vigor as might be affected by storm events, water temperature fluctuations, or winter ice scouring.

**Figure 5.**
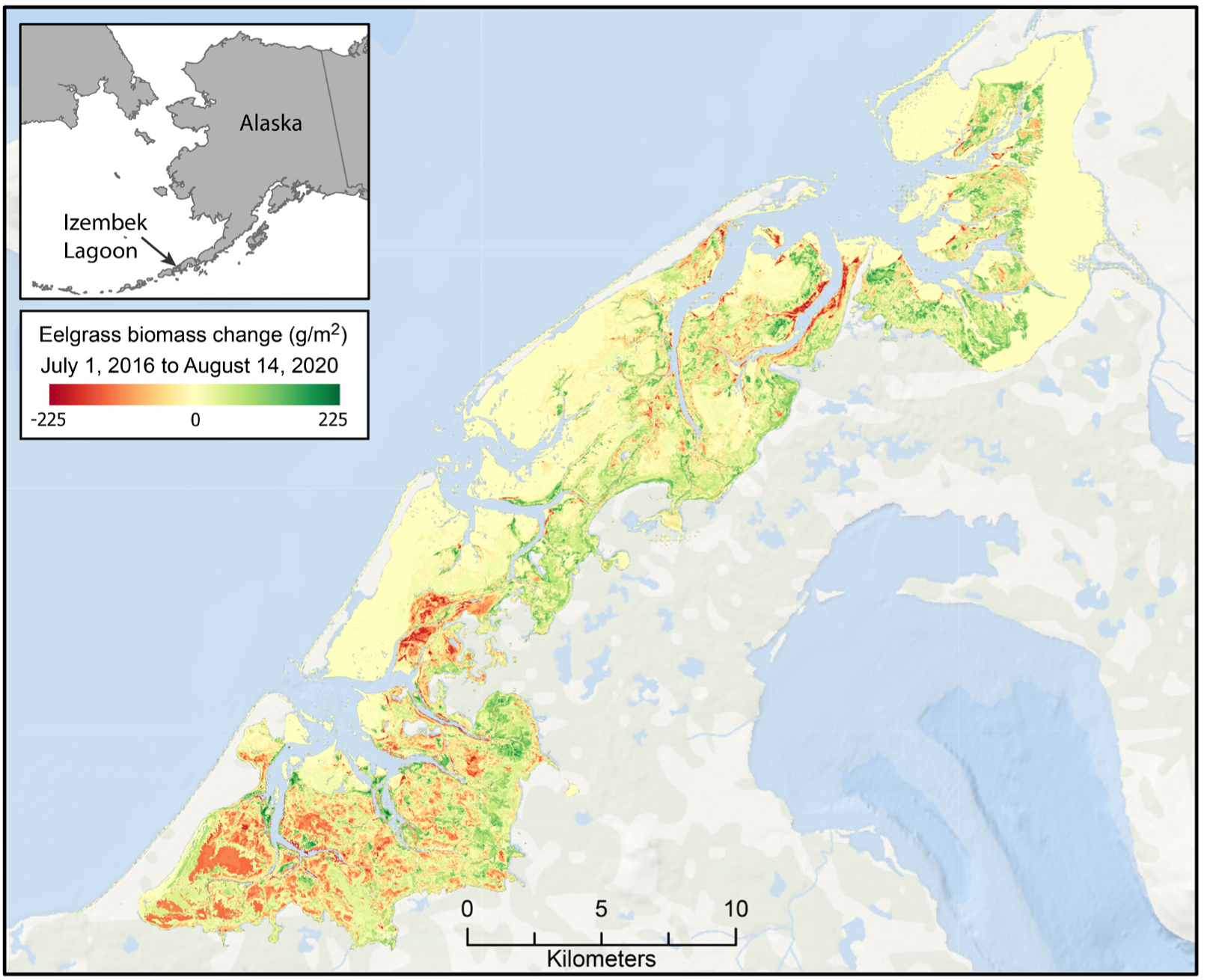
Estimated change in eelgrass biomass (g/m^2^) from July 1, 2016, to August 14, 2020, at Izembek Lagoon, Alaska.

We multiplied the biomass estimate for each spectral class in both Sentinel-2 maps (as mapped in Figure 3) by the respective class’s aerial extent, then summed across all classes to derive an estimate of the total eelgrass biomass throughout Izembek Lagoon on each respective image date (Table 6). The same methods used to attribute spectral classes by intersecting the plot estimates of biomass and assigning the spectral classes with median values after outliers were removed, were similarly applied using each ground plot’s upper (97.5%) and lower (2.5%) bounds of the 95% biomass credible interval (Table 6). The modest net increase in lagoon-wide biomass from 2016 to 2020 (Table 6) was inconclusive considering the broad 95% credible intervals of the annual point estimates.

**Table 6:**
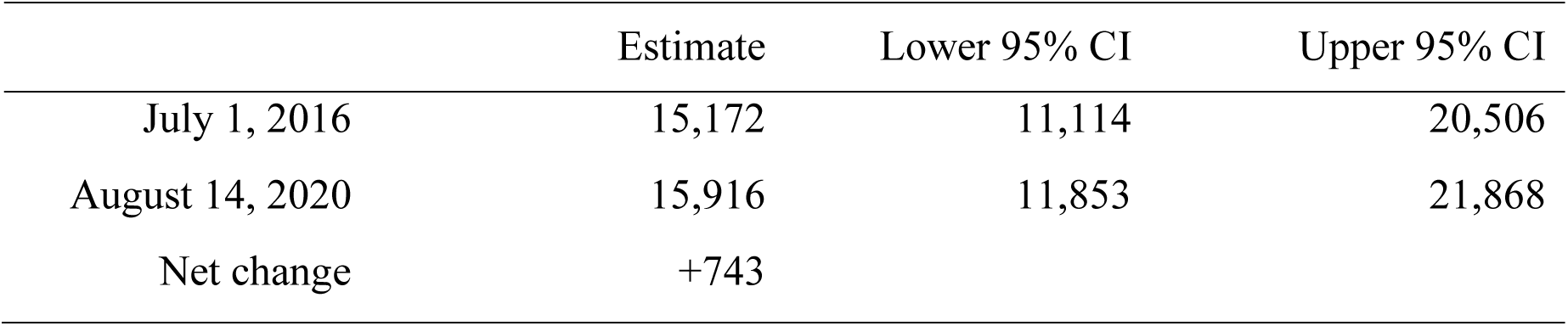
Estimated total eelgrass biomass (metric tons) in Izembek Lagoon, Alaska, on July 1, 2016, and on August 14, 2020. Lower and upper 95% CI are the upper and lower bounds of 95% Bayesian credible intervals.

### Data Availability

The maps of spectral classes, eelgrass attributes, and derivative maps of eelgrass metrics in geoTIFF and CSV format are available in Douglas et al. (2024, doi:10.5066/P1HLTAHD)

## Acknowledgments

We thank the European Space Agency and the Copernicus Data Space Ecosystem for making Sentinel-2 data available to the scientific community (https://dataspace.copernicus.eu). We acknowledge the use of imagery from the NASA Worldview application (https://worldview.earthdata.nasa.gov), part of the NASA Earth Science Data and Information System (ESDIS). This information product has been peer reviewed and approved for publication as a preprint by the U.S. Geological Survey. Any use of trade, firm or product names is for descriptive purposes only and does not imply endorsement by the U.S. Government.

## Appendices

**Figure A1.**
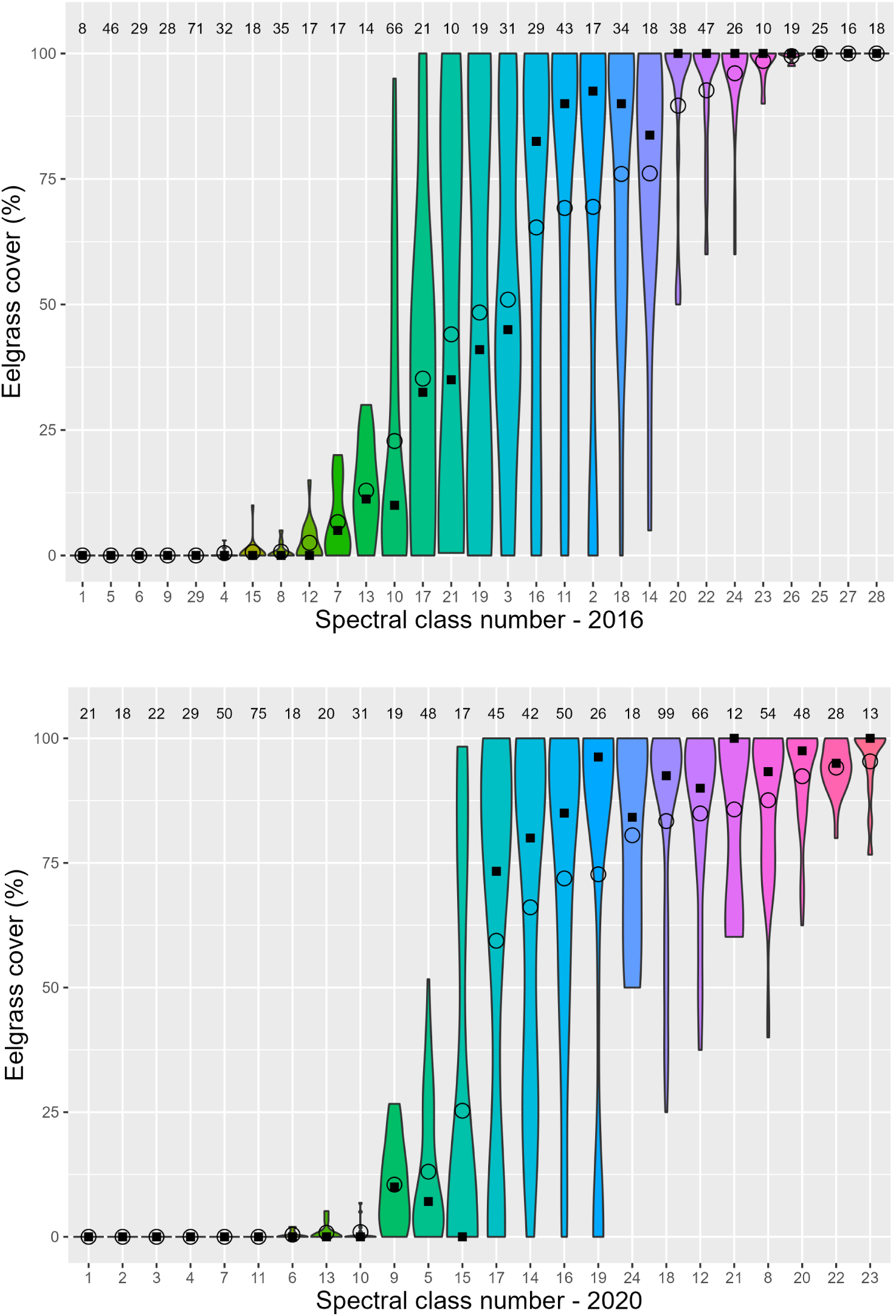
Eelgrass percent cover within each of the 29 spectral classes derived from the 2016 Sentinel-2 image (top) and the 24 spectral classes derived from the 2020 Sentinel-2 image (bottom). Class means are shown with an open circle, medians with a solid square. Spectral classes are sorted by ascending mean percent eelgrass cover. Violin shape depicts the frequency distribution of observations, scaled to constant width across classes.

**Figure A2.**
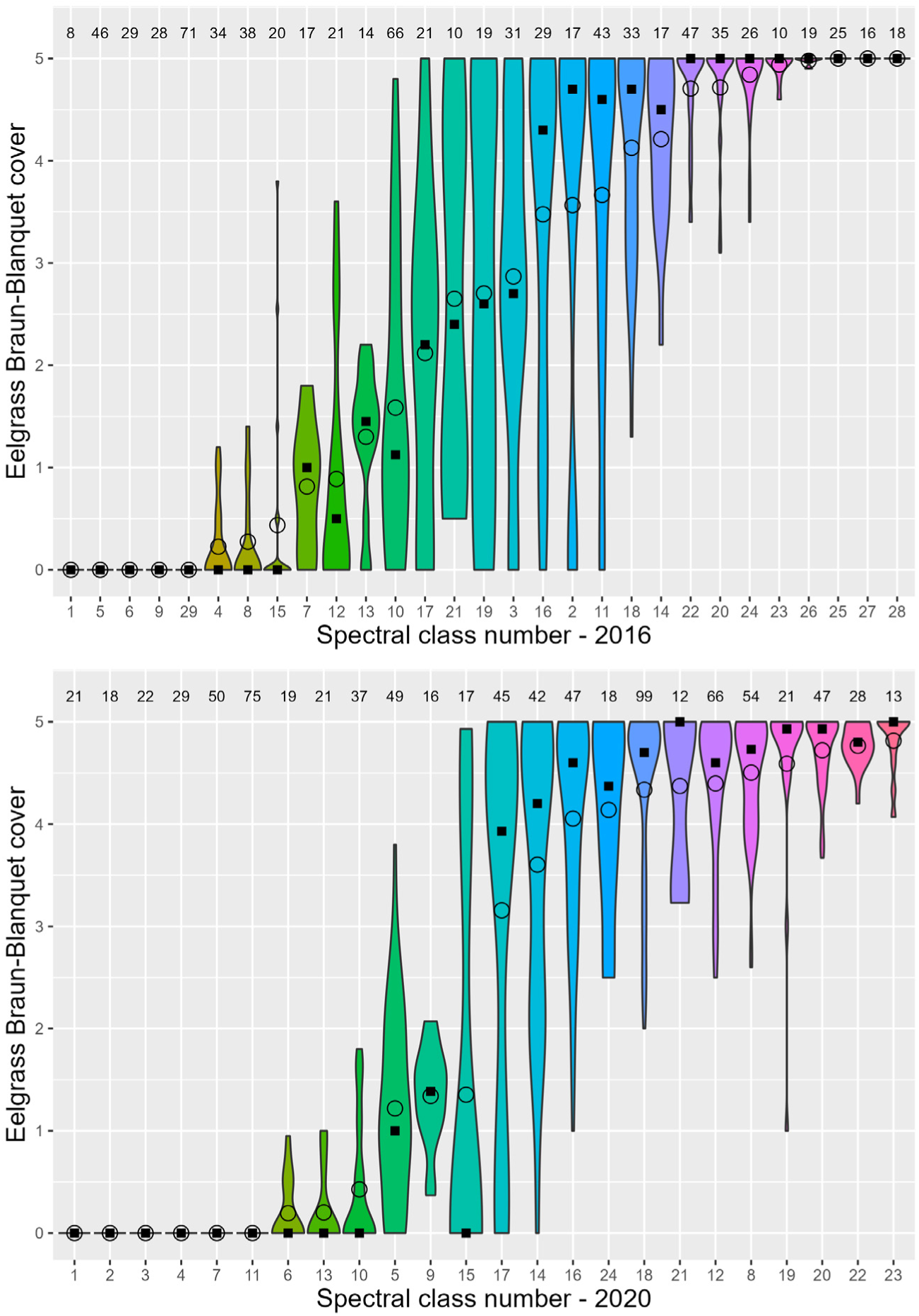
Eelgrass Braun-Blanquet cover within each of the 29 spectral classes from the 2016 Sentinel-2 image (top) and the 24 spectral classes derived from the 2020 Sentinel-2 image (bottom). Class means are shown with an open circle, medians with a solid square. Spectral classes are sorted by ascending mean Braun-Blanquet eelgrass cover. Violin shape depicts the frequency distribution of observations, scaled to constant width across classes.

**Figure A3.**
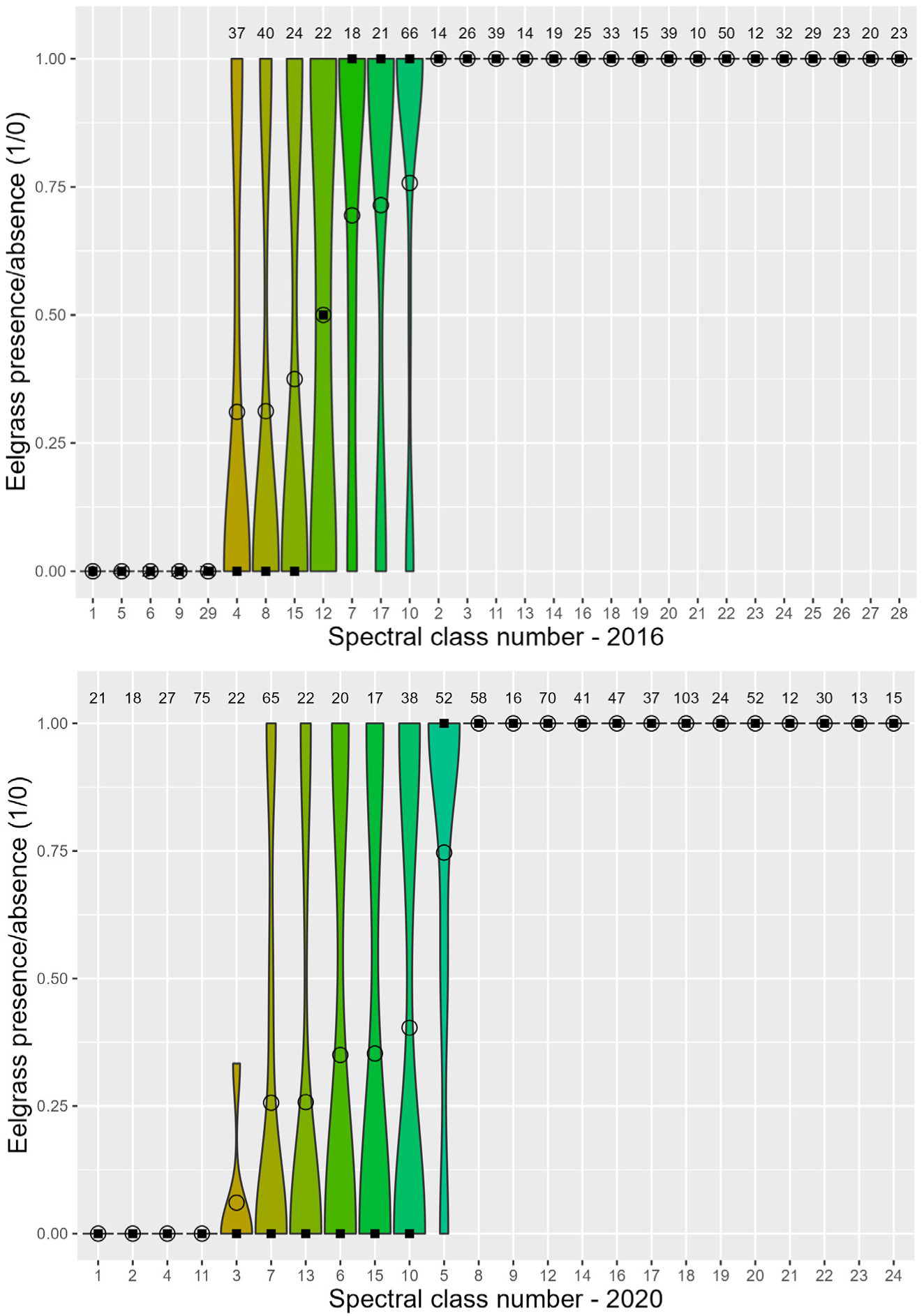
Eelgrass presence (1) or absence (0) within each of the 29 spectral classes derived from the 2016 Sentinel-2 image (top) and the 24 spectral classes derived from the 2020 Sentinel-2 image (bottom). Class means are shown with an open circle, medians with a solid square. Spectral classes are sorted by ascending mean. Sample sizes are shown across the top. Violin shape depicts the frequency distribution of observations, scaled to constant width across classes.

**Figure A4.**
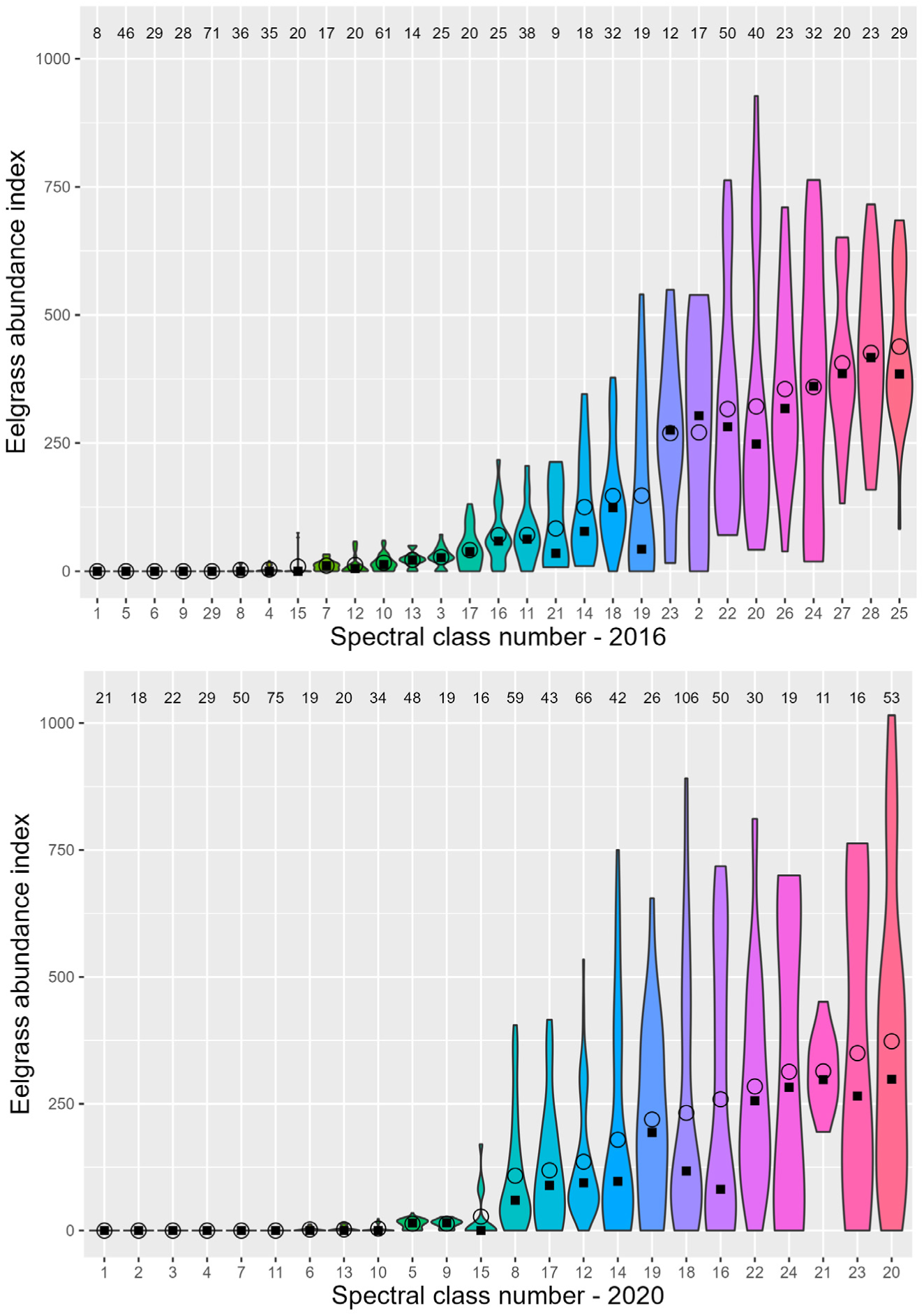
Eelgrass abundance index within each of the 29 spectral classes derived from the 2016 Sentinel-2 image (top) and the 24 spectral classes derived from the 2020 Sentinel-2 image (bottom). Class means are shown with an open circle, medians with a solid square. Spectral classes are sorted by ascending mean sample sizes are shown across the top. Violin shape depicts the frequency distribution of observations, scaled to a constant width across classes.

**Figure A5.**
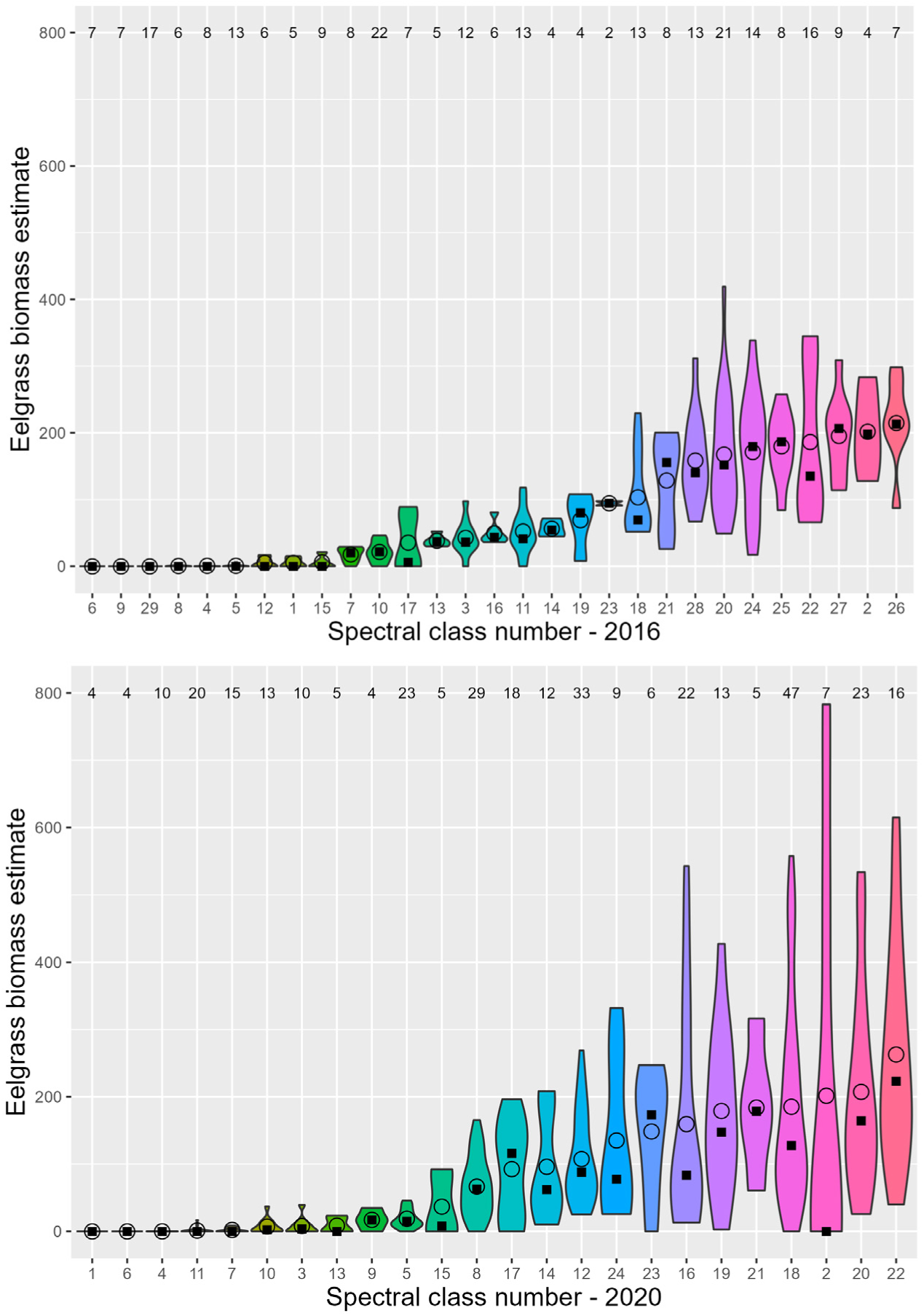
Eelgrass biomass estimates (g/m^2^) within each of the 29 spectral classes derived from the 2016 Sentinel-2 image (top) and the 24 spectral classes derived from the 2020 Sentinel-2 image (bottom). Class means are shown with an open circle, medians with a solid square. Spectral classes are sorted by ascending mean sample sizes are shown across the top. Violin shape depicts the frequency distribution of observations, scaled to a constant width across classes.

**Figure A6.**
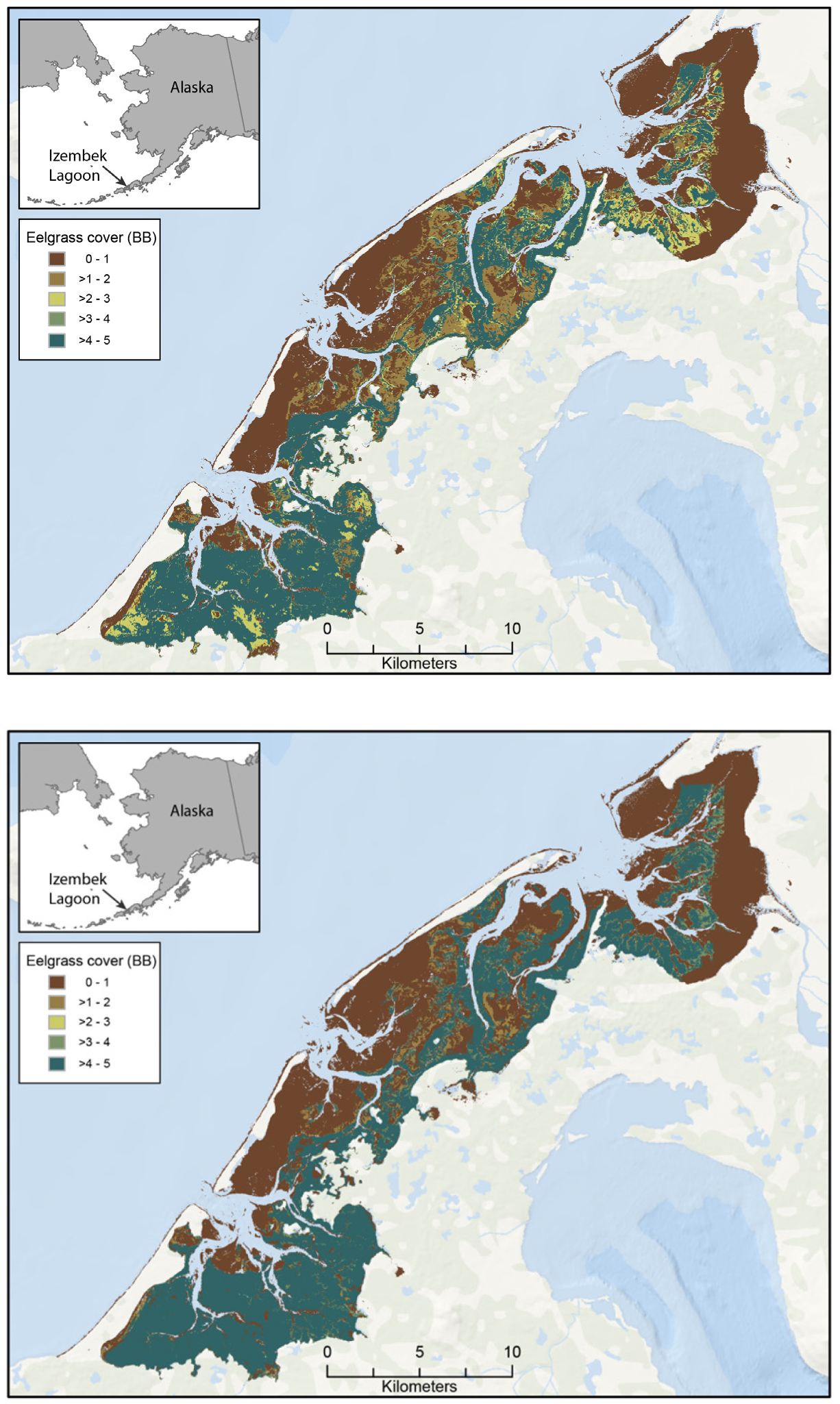
Maps of median eelgrass Braun-Blanquet cover at Izembek Lagoon, Alaska, derived from a Sentinel-2 satellite image collected during low tide on July 1, 2016 (top) and on August 14, 2020 (bottom). Twenty-nine/24 spectral classes were generated from the 2016/2020 Sentinel-2 multispectral images within the lagoon. In these maps, each spectral class has been color- shaded based on the median Braun-Blanquet cover category that was recorded at field plots located within the respective classes.

**Table A1.** Eelgrass percent cover statistics for each 2020 Sentinel-2 spectral class. IQR = interquartile range.

| *Class* | *N* | *Mean* | *Std dev* | *Median* | *IQR* |
| --- | --- | --- | --- | --- | --- |
| 1 | 21 | 0.00 | 0.00 | 0.00 | 0.00 |
| 2 | 18 | 0.00 | 0.00 | 0.00 | 0.00 |
| 3 | 22 | 0.00 | 0.00 | 0.00 | 0.00 |
| 4 | 29 | 0.00 | 0.00 | 0.00 | 0.00 |
| 7 | 50 | 0.00 | 0.00 | 0.00 | 0.00 |
| 11 | 75 | 0.00 | 0.00 | 0.00 | 0.00 |
| 6 | 18 | 0.43 | 0.76 | 0.00 | 0.75 |
| 13 | 20 | 0.82 | 1.75 | 0.00 | 0.00 |
| 10 | 31 | 0.95 | 2.02 | 0.00 | 0.55 |
| 9 | 19 | 10.47 | 8.58 | 10.00 | 11.84 |
| 5 | 48 | 13.07 | 13.83 | 7.08 | 23.50 |
| 15 | 17 | 25.29 | 40.21 | 0.00 | 66.67 |
| 17 | 45 | 59.39 | 40.07 | 73.33 | 90.00 |
| 14 | 42 | 66.10 | 33.87 | 80.00 | 65.83 |
| 16 | 50 | 71.90 | 31.12 | 85.00 | 44.79 |
| 19 | 26 | 72.72 | 41.33 | 96.25 | 41.25 |
| 24 | 18 | 80.56 | 19.84 | 84.17 | 34.38 |
| 18 | 99 | 83.40 | 21.31 | 92.50 | 20.00 |
| 12 | 66 | 84.93 | 17.30 | 90.00 | 18.33 |
| 21 | 12 | 85.75 | 18.27 | 100.00 | 36.50 |
| 8 | 54 | 87.59 | 14.27 | 93.33 | 21.25 |
| 20 | 48 | 92.40 | 9.94 | 97.50 | 10.83 |
| 22 | 28 | 94.14 | 4.98 | 95.00 | 6.87 |
| 23 | 13 | 95.38 | 7.61 | 100.00 | 5.00 |

*Spectral classes sorted by mean percent cover*

**Table A2.** Eelgrass Braun-Blanquet cover category statistics for each 2020 Sentinel-2 spectral class. IQR = interquartile range.

| *Class* | *N* | *Mean* | *Std dev* | *Median* | *IQR* |
| --- | --- | --- | --- | --- | --- |
| 1 | 21 | 0.00 | 0.00 | 0.00 | 0.00 |
| 2 | 18 | 0.00 | 0.00 | 0.00 | 0.00 |
| 3 | 22 | 0.00 | 0.00 | 0.00 | 0.00 |
| 4 | 29 | 0.00 | 0.00 | 0.00 | 0.00 |
| 7 | 50 | 0.00 | 0.00 | 0.00 | 0.00 |
| 11 | 75 | 0.00 | 0.00 | 0.00 | 0.00 |
| 6 | 19 | 0.19 | 0.31 | 0.00 | 0.50 |
| 13 | 21 | 0.20 | 0.38 | 0.00 | 0.00 |
| 10 | 37 | 0.43 | 0.62 | 0.00 | 0.57 |
| 5 | 49 | 1.22 | 0.93 | 1.00 | 1.37 |
| 9 | 16 | 1.34 | 0.48 | 1.38 | 0.50 |
| 15 | 17 | 1.35 | 2.06 | 0.00 | 3.67 |
| 17 | 45 | 3.16 | 1.93 | 3.93 | 3.80 |
| 14 | 42 | 3.60 | 1.44 | 4.20 | 2.65 |
| 16 | 47 | 4.05 | 1.05 | 4.60 | 1.46 |
| 24 | 18 | 4.14 | 0.94 | 4.37 | 1.38 |
| 18 | 99 | 4.34 | 0.85 | 4.70 | 0.80 |
| 21 | 12 | 4.37 | 0.81 | 5.00 | 1.63 |
| 12 | 66 | 4.40 | 0.69 | 4.60 | 0.73 |
| 8 | 54 | 4.50 | 0.57 | 4.73 | 0.85 |
| 19 | 21 | 4.59 | 0.94 | 4.93 | 0.40 |
| 20 | 47 | 4.72 | 0.36 | 4.93 | 0.40 |
| 22 | 28 | 4.77 | 0.20 | 4.80 | 0.27 |
| 23 | 13 | 4.82 | 0.30 | 5.00 | 0.20 |

*Spectral classes sorted by mean BB cover category*

**Table A3.** Eelgrass presence (binary) statistics for each 2020 Sentinel-2 spectral class. IQR = interquartile range.

| *Class* | *N* | *Mean* | *Std dev* | *Median* | *IQR* |
| --- | --- | --- | --- | --- | --- |
| 1 | 21 | 0.00 | 0.00 | 0 | 0.00 |
| 2 | 18 | 0.00 | 0.00 | 0 | 0.00 |
| 4 | 27 | 0.00 | 0.00 | 0 | 0.00 |
| 11 | 75 | 0.00 | 0.00 | 0 | 0.00 |
| 3 | 22 | 0.06 | 0.13 | 0 | 0.00 |
| 7 | 65 | 0.26 | 0.41 | 0 | 0.50 |
| 13 | 22 | 0.26 | 0.44 | 0 | 0.50 |
| 6 | 20 | 0.35 | 0.49 | 0 | 1.00 |
| 15 | 17 | 0.35 | 0.49 | 0 | 1.00 |
| 10 | 38 | 0.40 | 0.48 | 0 | 1.00 |
| 5 | 52 | 0.75 | 0.39 | 1 | 0.50 |
| 8 | 58 | 1.00 | 0.00 | 1 | 0.00 |
| 9 | 16 | 1.00 | 0.00 | 1 | 0.00 |
| 12 | 70 | 1.00 | 0.00 | 1 | 0.00 |
| 14 | 41 | 1.00 | 0.00 | 1 | 0.00 |
| 16 | 47 | 1.00 | 0.00 | 1 | 0.00 |
| 17 | 37 | 1.00 | 0.00 | 1 | 0.00 |
| 18 | 103 | 1.00 | 0.00 | 1 | 0.00 |
| 19 | 24 | 1.00 | 0.00 | 1 | 0.00 |
| 20 | 52 | 1.00 | 0.00 | 1 | 0.00 |
| 21 | 12 | 1.00 | 0.00 | 1 | 0.00 |
| 22 | 30 | 1.00 | 0.00 | 1 | 0.00 |
| 23 | 13 | 1.00 | 0.00 | 1 | 0.00 |
| 24 | 15 | 1.00 | 0.00 | 1 | 0.00 |

*Spectral classes sorted by mean presence*

**Table A4.**
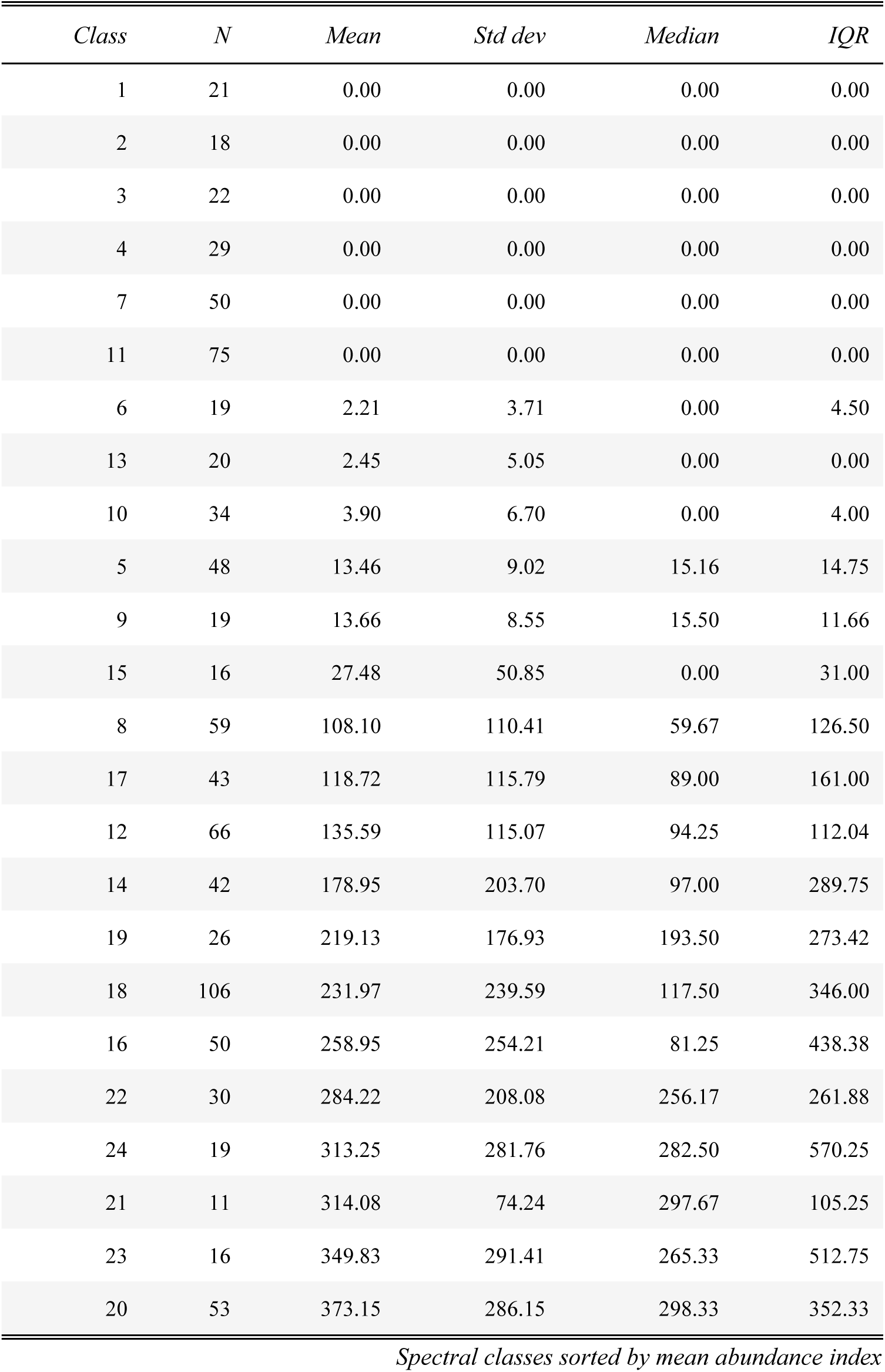
Eelgrass abundance index statistics for each 2020 Sentinel-2 spectral class. IQR = interquartile range.

**Table A5.** Eelgrass percent cover statistics for each 2016 Sentinel-2 spectral class.

| *Class* | *N* | *Mean* | *Std dev* | *Median* | *IOR* |
| --- | --- | --- | --- | --- | --- |
| 1 | 8 | 0.00 | 0.00 | 0.00 | 0.00 |
| 5 | 46 | 0.00 | 0.00 | 0.00 | 0.00 |
| 6 | 29 | 0.00 | 0.00 | 0.00 | 0.00 |
| 9 | 28 | 0.00 | 0.00 | 0.00 | 0.00 |
| 29 | 71 | 0.00 | 0.00 | 0.00 | 0.00 |
| 4 | 32 | 0.47 | 0.95 | 0.00 | 0.12 |
| 15 | 18 | 0.67 | 2.35 | 0.00 | 0.00 |
| 8 | 35 | 0.76 | 1.57 | 0.00 | 0.00 |
| 12 | 17 | 2.53 | 3.93 | 0.00 | 5.00 |
| 7 | 17 | 6.65 | 7.33 | 5.00 | 14.50 |
| 13 | 14 | 12.91 | 9.68 | 11.25 | 11.88 |
| 10 | 66 | 22.80 | 27.92 | 10.00 | 37.50 |
| 17 | 21 | 35.24 | 32.00 | 32.50 | 55.00 |
| 21 | 10 | 44.05 | 42.42 | 35.00 | 75.00 |
| 19 | 19 | 48.39 | 41.91 | 41.00 | 87.25 |
| 3 | 31 | 50.97 | 34.34 | 45.00 | 56.25 |
| 16 | 29 | 65.34 | 37.48 | 82.50 | 60.00 |
| 11 | 43 | 69.22 | 36.32 | 90.00 | 55.00 |
| 2 | 17 | 69.44 | 41.54 | 92.50 | 47.50 |
| 18 | 34 | 76.00 | 28.25 | 90.00 | 40.00 |
| 14 | 18 | 76.11 | 26.31 | 83.75 | 37.50 |
| 20 | 38 | 89.61 | 17.32 | 100.00 | 10.00 |
| 22 | 47 | 92.66 | 12.11 | 100.00 | 10.00 |
| 24 | 26 | 96.06 | 9.90 | 100.00 | 0.00 |
| 23 | 10 | 98.50 | 3.37 | 100.00 | 0.00 |
| 26 | 19 | 99.47 | 1.05 | 100.00 | 0.00 |
| 25 | 25 | 100.00 | 0.00 | 100.00 | 0.00 |
| 27 | 16 | 100.00 | 0.00 | 100.00 | 0.00 |
| 28 | 18 | 100.00 | 0.00 | 100.00 | 0.00 |

*Spectral classes sorted by mean percent cover*

**Table A6.**
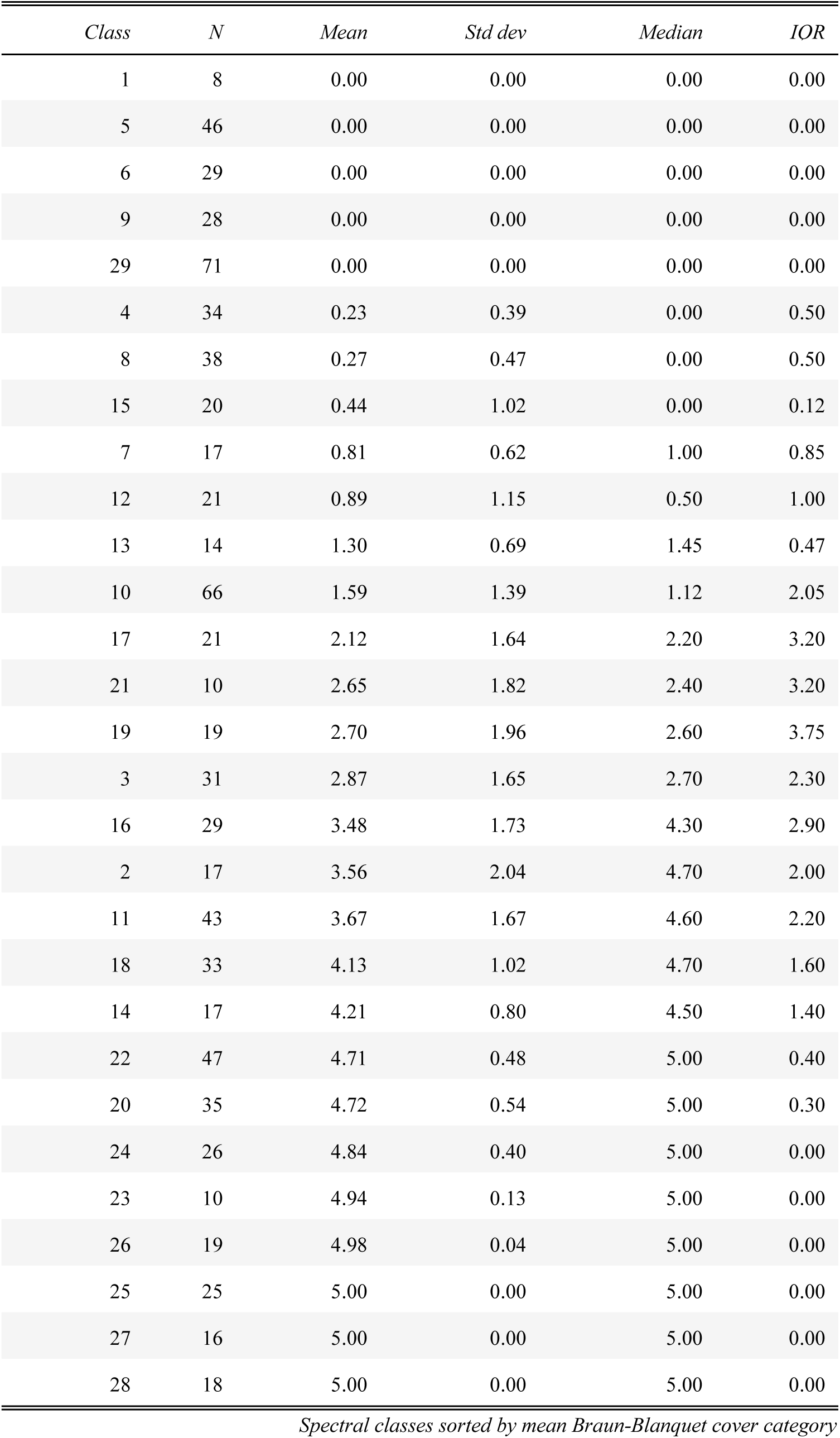
Eelgrass Braun-Blanquet cover category statistics for each 2016 Sentinel-2.

**Table A7.**
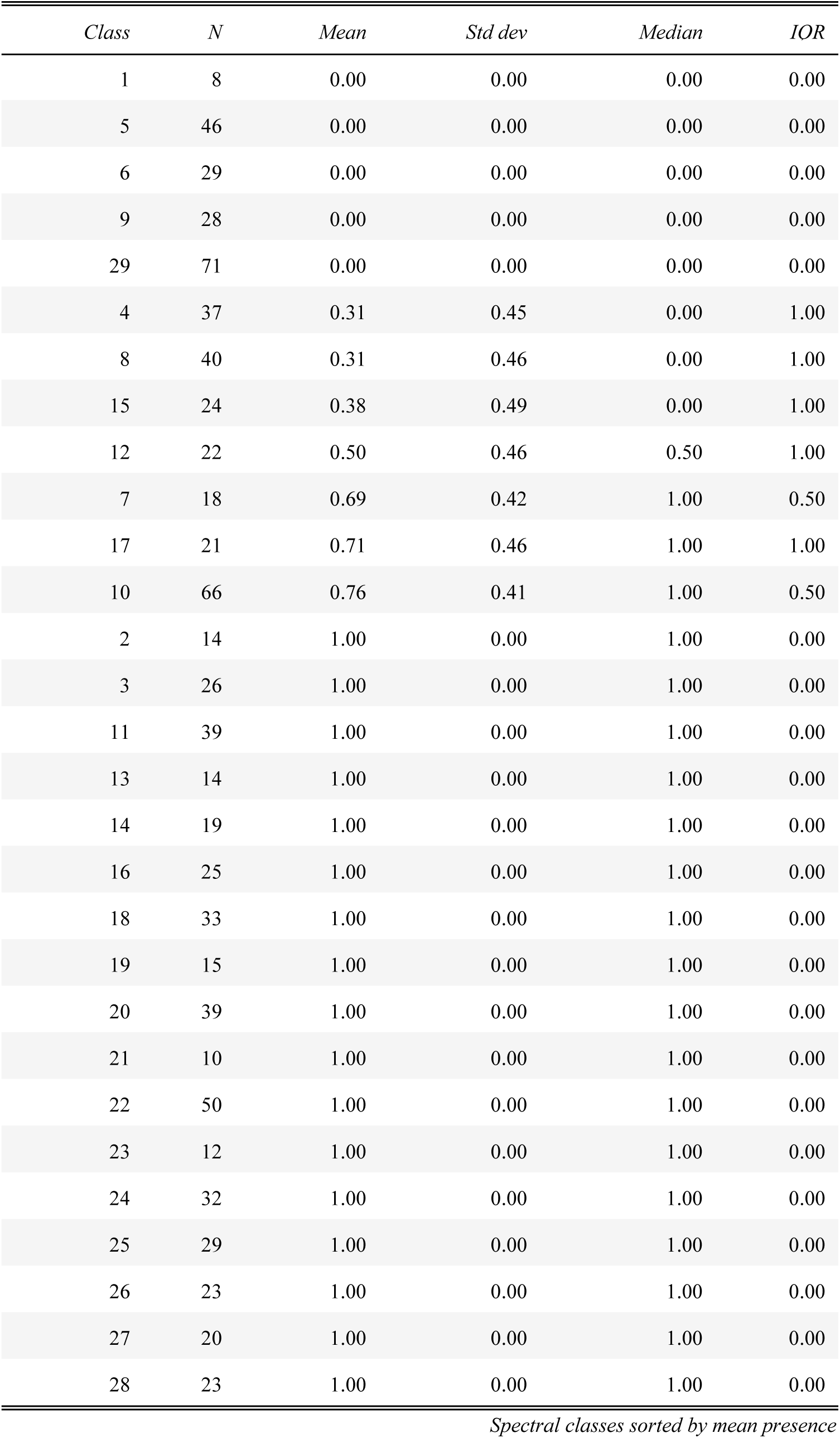
Eelgrass presence (binary) statistics for each 2016 Sentinel-2 spectral class.

**Table A8.** Eelgrass abundance index statistics for each 2016 Sentinel-2 spectral class.

| *Class* | *N* | *Mean* | *Std dev* | *Median* | *IOR* |
| --- | --- | --- | --- | --- | --- |
| 1 | 8 | 0.00 | 0.00 | 0.00 | 0.00 |
| 5 | 46 | 0.00 | 0.00 | 0.00 | 0.00 |
| 6 | 29 | 0.00 | 0.00 | 0.00 | 0.00 |
| 9 | 28 | 0.00 | 0.00 | 0.00 | 0.00 |
| 29 | 71 | 0.00 | 0.00 | 0.00 | 0.00 |
| 8 | 36 | 2.58 | 4.94 | 0.00 | 1.25 |
| 4 | 35 | 3.63 | 5.99 | 0.00 | 6.00 |
| 15 | 20 | 8.95 | 21.83 | 0.00 | 1.75 |
| 7 | 17 | 10.94 | 9.86 | 11.00 | 11.50 |
| 12 | 20 | 12.12 | 18.12 | 5.25 | 12.25 |
| 10 | 61 | 16.48 | 14.97 | 13.00 | 17.00 |
| 13 | 14 | 22.21 | 14.76 | 22.00 | 11.00 |
| 3 | 25 | 27.56 | 18.59 | 26.50 | 17.00 |
| 17 | 20 | 41.27 | 39.07 | 38.00 | 50.75 |
| 16 | 25 | 70.26 | 54.88 | 59.00 | 41.00 |
| 11 | 38 | 70.95 | 52.66 | 62.50 | 57.50 |
| 21 | 9 | 83.56 | 82.25 | 35.00 | 144.00 |
| 14 | 18 | 125.06 | 102.46 | 78.00 | 151.62 |
| 18 | 32 | 146.75 | 100.89 | 124.25 | 115.38 |
| 19 | 19 | 147.71 | 168.52 | 43.00 | 224.50 |
| 23 | 12 | 269.21 | 156.44 | 275.25 | 168.88 |
| 2 | 17 | 270.74 | 194.29 | 303.50 | 310.00 |
| 22 | 50 | 316.74 | 207.08 | 281.75 | 275.50 |
| 20 | 40 | 321.70 | 252.36 | 248.00 | 288.88 |
| 26 | 23 | 355.57 | 172.81 | 317.50 | 245.00 |
| 24 | 32 | 359.61 | 242.92 | 361.00 | 484.25 |
| 27 | 20 | 405.68 | 147.57 | 386.00 | 152.50 |
| 28 | 23 | 425.98 | 153.71 | 417.00 | 207.00 |
| 25 | 29 | 438.30 | 140.46 | 385.00 | 233.00 |

*Spectral classes sorted by mean abundance index*

**Table A9.**
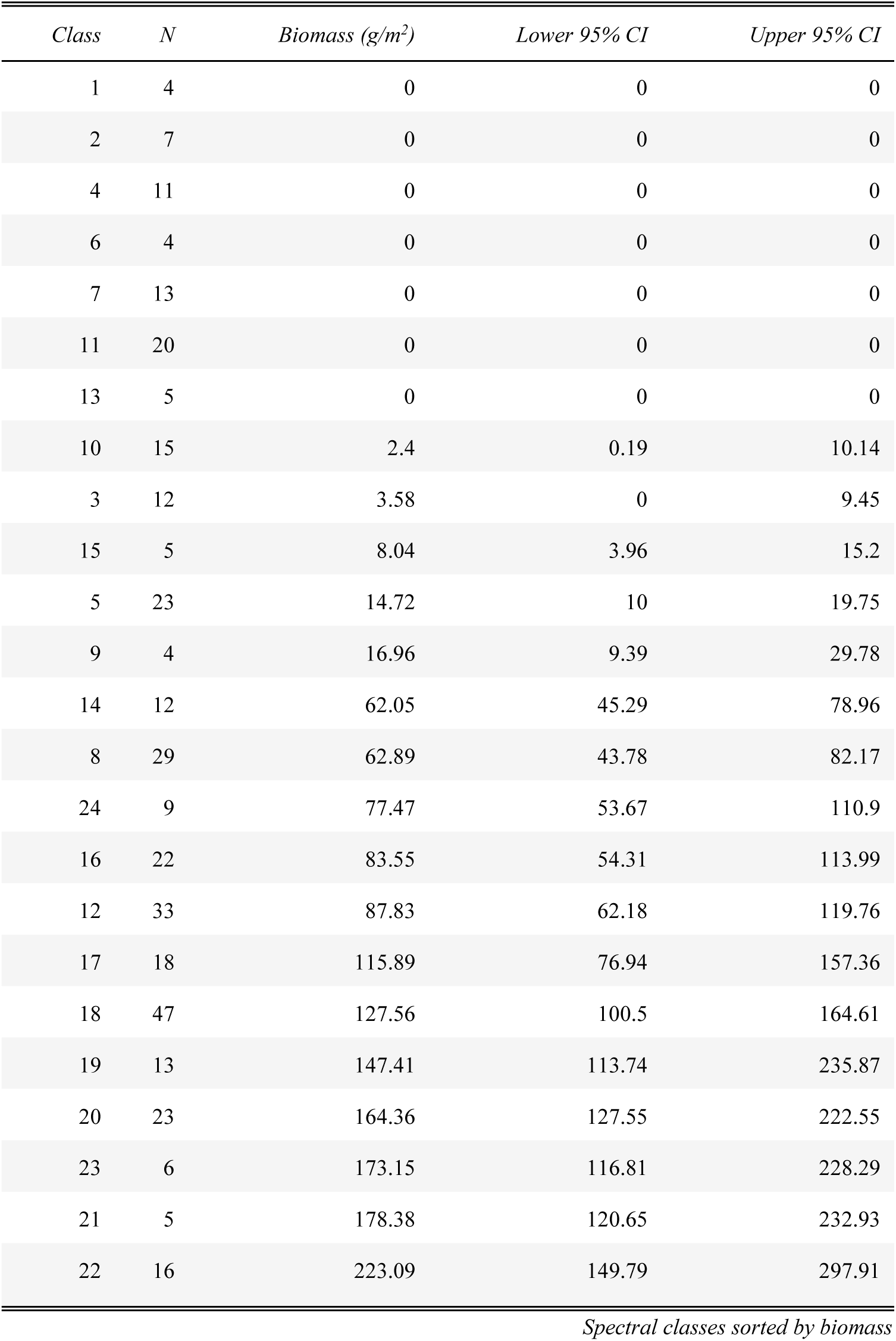
Median eelgrass biomass estimates for each 2020 Sentinel-2 spectral class.

| *Class* | *N* | *Biomass (g/m<sup>2</sup>)* | *Lower 95% CI* | *Upper 95% CI* |
| --- | --- | --- | --- | --- |
| 1 | 4 | 0 | 0 | 0 |
| 2 | 7 | 0 | 0 | 0 |
| 4 | 11 | 0 | 0 | 0 |
| 6 | 4 | 0 | 0 | 0 |
| 7 | 13 | 0 | 0 | 0 |
| 11 | 20 | 0 | 0 | 0 |
| 13 | 5 | 0 | 0 | 0 |
| 10 | 15 | 2.4 | 0.19 | 10.14 |
| 3 | 12 | 3.58 | 0 | 9.45 |
| 15 | 5 | 8.04 | 3.96 | 15.2 |
| 5 | 23 | 14.72 | 10 | 19.75 |
| 9 | 4 | 16.96 | 9.39 | 29.78 |
| 14 | 12 | 62.05 | 45.29 | 78.96 |
| 8 | 29 | 62.89 | 43.78 | 82.17 |
| 24 | 9 | 77.47 | 53.67 | 110.9 |
| 16 | 22 | 83.55 | 54.31 | 113.99 |
| 12 | 33 | 87.83 | 62.18 | 119.76 |
| 17 | 18 | 115.89 | 76.94 | 157.36 |
| 18 | 47 | 127.56 | 100.5 | 164.61 |
| 19 | 13 | 147.41 | 113.74 | 235.87 |
| 20 | 23 | 164.36 | 127.55 | 222.55 |
| 23 | 6 | 173.15 | 116.81 | 228.29 |
| 21 | 5 | 178.38 | 120.65 | 232.93 |
| 22 | 16 | 223.09 | 149.79 | 297.91 |

*Spectral classes sorted by biomass*

**Table A10.**
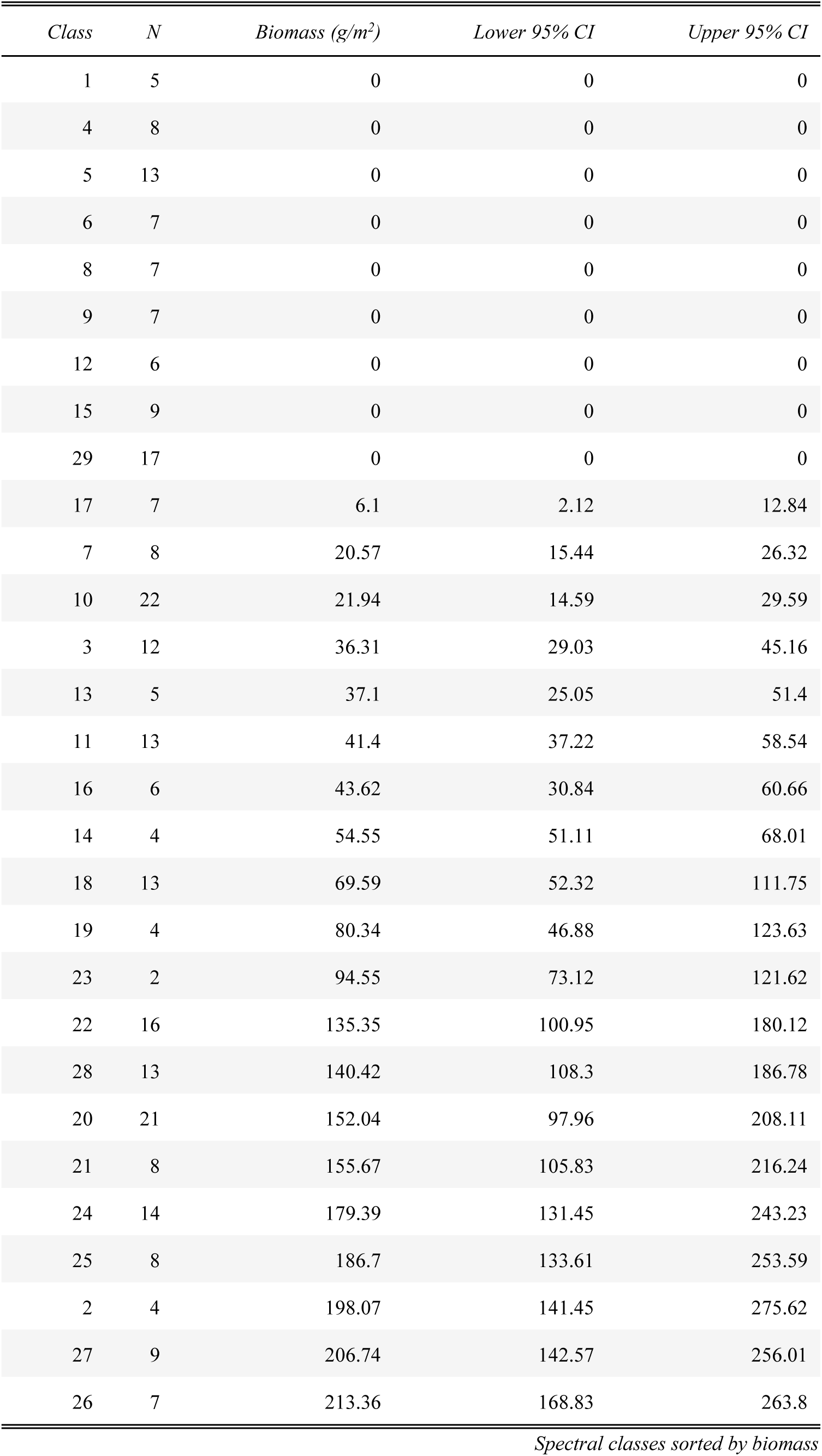
Median eelgrass biomass estimates for each 2016 Sentinel-2 spectral class.

## References

1. Aucoin, P.J., and J. Stewart, 1978, Earth Observations Division version of the Laboratory for Applications of Remote Sensing system (EOD-LARSYS) user guide for the IBM 370/148. Volume 1: System overview, JSC-13821, National Aeronautics Space Administration, Lyndon B. Johnson Space Center, Houston, TX. 227 pp. https://ntrs.nasa.gov/api/citations/19800018225/downloads/19800018225.pdf

2. Bartenfelder, A., W.J. Kenworthy, B. Puckett, C. Deaton, and J.C. Jarvis, 2022, The abundance and persistence of temperate and tropical seagrasses at their edge-of-range in the Western Atlantic Ocean. Frontiers Marine Science 9: 917237 10.3389/fmars.2022.917237

3. Braun-Blanquet, J., 1972. Plant sociology: the study of plant communities: authorized English translation of Pflanzensoziologie. (Translated, reviewed and edited by Fuller, G.D. and Conard, H.S., editors) New York: Hafner Publishing Co.

4. Brower Jr., W.A., R.G. Baldwin, L.N. Williams Jr., J.L Wise, and L.D. Leslie, 1988, Climatic atlas: the Outer Continental Shelf waters and coastal regions of Alaska, Vol 2. Arctic environmental information and data center, Univ. of Alaska, Anchorage, AK, pp. 519.

5. Douglas, D.C., M.D. Fleming, V.P. Patil, and D.H. Ward, 2024, Eelgrass (*Zostera marina*) maps from 2016 and 2020, at Izembek National Wildlife Refuge, Alaska: U.S. Geological Survey data release, 10.5066/P1HLTAHD

6. Drusch, M., U. Del Bello, S. Carlier, O. Colin, V. Fernandez, F. Gascon, B. Hoersch, C. Isola, P. Laberinti, and P. Martimort, 2012, Sentinel-2: ESA’s optical high-resolution mission for GMES operational services. Remote Sens. Environ. 120:25–36. 10.1016/j.rse.2011.11.026

7. Hogrefe, K.R., D.H. Ward, T.F. Donnelly, and N. Dau, 2014, Establishing a baseline for regional scale monitoring of eelgrass (Zostera marina) habitat on the Lower Alaska Peninsula. Remote Sensing 6(12):12447–12477. 10.3390/rs61212447

8. Li, J., and D.P. Roy, 2017, A global analysis of Sentinel-2A, Sentinel-2B and Landsat-8 data revisit intervals and implications for terrestrial monitoring, Remote Sensing 9(9):902. 10.3390/rs9090902

9. Loveland, T.R., and J.R. Irons, 2016, Landsat 8: The plans, the reality, and the legacy, Remote Sensing of Environment, 185:1–6 10.1016/j.rse.2016.07.033

10. McRoy, C.P., 1966, Standing stocks and ecology of eelgrass (Zostera marina) at Izembek Lagoon, Alaska, MS Thesis, Univ. of Washington, Seattle, WA, pp. 138

11. Munsch, S.H., F.L. Beaty, K.M. Beheshti, W.B. Chesney, and others, 2023, Northeast Pacific eelgrass dynamics: interannual expansion distances and meadow area variation over time. Marine Ecology Progress Series 705:61–75. 10.3354/meps14248

12. Neckles, H.A, B.S. Kopp, B.J. Peterson, and P.S. Pooler, 2012, Integrating scales of seagrass monitoring to meet conservation needs, Estuaries and Coasts, 35:23–46. 10.1007/s12237-011-9410-x

13. Overland, J.E., E. Siddon, G. Sheffield, T.J. Ballinger, and C. Szuwalski, 2024, Transformative ecological and human impacts from diminished sea ice in the northern Bering Sea, Weather, Climate, and Society, 16:303–313, 10.1175/WCAS-D-23-0029.1

14. Patil, V.P., 2024, Eelgrass biomass model (ver 1.0.1, May 2024): U.S. Geological Survey software release, 10.5066/P13EG9KS

15. Phiri, D., M. Simwanda, S. Salekin, V.R. Nyirenda, Y. Jurayama, and M. Ranagalage, 2020, Sentinel-2 data for land cover/use mapping: A review, Remote Sensing, 12(14), 2291, 10.3390/rs12142291

16. Ramsar, 2023, The list of wetlands of international importance, https://www.ramsar.org/sites/default/files/documents/library/sitelist.pdf

17. Sherman, K., and L.A. DeBruyckere, 2018, Eelgrass habitats on the U.S. west coast. State of the knowledge of eelgrass ecosystem services and eelgrass extent. A publication prepared by the Pacific Marine and Estuarine Fish Habitat Partnership for The Nature Conservancy. 67pp. https://www.pacificfishhabitat.org/wp-content/uploads/2017/09/EelGrass_Report_Final_ForPrint_web.pdf

18. Swain, P.H. and R.C. King, 1973, Two effective feature selection criteria for multispectral remote sensing. LARS Technical Reports. Paper 39. http://docs.lib.purdue.edu/larstech/39

19. Ward, D.H., C.J. Markon, and D.C. Douglas, 1997, Distribution and stability of eelgrass beds at Izembek Lagoon, Alaska, Aquatic Botany, 58:229–240. 10.1016/S0304-3770(97)00037-5

20. Ward, D.H., A. Morton, T.L. Tibbitts, D.C. Douglas, and E. Carrera-Gonzalez, 2003, Long-term change in eelgrass distribution at Bahia San Quintin, Baja California, Mexico, using satellite imagery, Estuaries, 26:1529–1539. 10.1007/BF02803661

21. Ward, D.H., and C.L. Amundson, 2019, Monitoring annual trends in abundance of eelgrass (Zostera marina) at Izembek National Wildlife Refuge, Alaska, 2018, U.S. Geological Survey Open-File Report 2019-1042, 8 p., 10.3133/ofr20191042.

22. Ward, D. H., 2021, Point sampling data for eelgrass (Zostera marina) and seaweed distribution and abundance in bays adjacent to the Izembek National Wildlife Refuge, Alaska (ver 4.0, June 2024): U.S. Geological Survey data release, 10.5066/P9ZUDIOH

23. Ward, D.H., K.R. Hogrefe, T.F. Donnelly, L.L. Fairchild, K.M. Sowl, and S.C. Lindstrom, 2022, Abundance and distribution of eelgrass (Zostera marina) and seaweeds at Izembek National Wildlife Refuge, Alaska, 2007–10: U.S. Geological Survey Open-File Report 2020–1035, 30 p., 10.3133/ofr20201035.

24. Wood, K.R., N.A. Bond, S.L. Danielson, J.E. Overland, S.A. Salo, P.J. Stabeno, and J. Whitefield, 2015, A decade of environmental change in the Pacific Arctic region, Prog. in Oceanography, 136:12–31, 10.1016/j.pocean.2015.05.005

